# Cyclic di-AMP signalling affects cell division and penicillin susceptibility in *Streptococcus pneumoniae*

**DOI:** 10.64898/2026.09.06.749708

**Authors:** Ragnhild Sødal Gjennestad, Silje Henriette Aadne, Morten Kjos, Daniel Straume

**Affiliations:** Faculty of Chemistry, Biotechnology and Food Science, Norwegian University of Life Sciences, Norway

## Abstract

Resistance to β-lactam antibiotics in the human pathogen *Streptococcus pneumoniae* is mainly attributed to the acquisition of mutated versions of *pbp2x*, *pbp2b* and *pbp1a*, encoding functional penicillin binding proteins with reduced affinity to β-lactams. This enables the bacterium to synthesise peptidoglycan required for cell division in the presence of higher β-lactam concentrations. In addition, it has recently been shown that increased levels of the second messenger molecule cyclic di-AMP, resulting from mutations that compromise the cyclic di-AMP degrading phosphodiesterase Pde1, contribute to decreased β-lactam susceptibility in this species. A link between cyclic di-AMP levels and β-lactam resistance has also been reported in several other Gram-positive bacteria. Here, we further investigated this link in *S. pneumoniae* and present evidence supporting that cyclic di-AMP could be involved in regulation of cell division. We confirm that elevated cyclic di-AMP levels decreased penicillin susceptibility and cell size in *pde1* mutants. Next, we show that low cyclic di-AMP levels had the opposite effects in which cells became more susceptible to penicillin and displayed an elongated morphology with multiple incomplete septa. Using fluorescence microscopy and bacterial two-hybrid assays, we found that proteins involved in cyclic di-AMP synthesis (CdaA, CdaR) and degradation (Pde1) were enriched around the division site and that they most probably interact with each other. Finally, we identified changes to the cell wall stem peptide composition of Δ*pde1* cells. Together our findings indicate that cyclic di-AMP could function as a regulator of the division machinery.

## Introduction

*Streptococcus pneumoniae*, also known as the pneumococcus, is a human pathogen that can cause a wide range of diseases in the respiratory system including sinusitis, otitis media and pneumonia as well as systemic infections leading to bacteriemia and meningitis (Henriques-Normark & Tuomanen, 2013; Kadioglu et al., 2008; Narciso et al., 2025). Annually, this pathogen causes about 1 million deaths and is the leading cause of infectious death in children and elderly (Troeger et al., 2018). β-lactam antibiotics, such as penicillins, have commonly been used for treating pneumococcal infections. However, the number of infections caused by β-lactam resistant isolates is increasing making treatment more difficult, which results in increased use of other broad-spectrum antibiotics and last resort drugs (Gergova et al., 2024; Li et al., 2022).

In *S. pneumoniae*, β-lactam resistance is mainly caused by the strains acquiring mutated versions of its targets, i.e. the penicillin binding proteins (PBPs) which are enzymes critical for synthesis of peptidoglycan in the bacterial cell wall (Hakenbeck et al., 2012; Hakenbeck et al., 1980; Sauvage et al., 2008; Zighelboim & Tomasz, 1980). PBPs synthesise peptidoglycan by performing glycosyl transferase and/or transpeptidase reactions. Class A PBPs perform both reactions while Class B PBPs only perform transpeptidation. Class A PBPs form glycan chains by polymerising units of the disaccharide *N*-acetylglucosamine (Glc*N*Ac) and N-acetylmuramic acid (Mur*N*Ac) (Sauvage et al., 2008). A pentapeptide composed of L-Ala-D-iGln-L-Lys-D-Ala-D-Ala (in *S. pneumoniae*) is attached to Mur*N*Ac and used by PBPs to crosslink adjacent glycan chains by transpeptidation reactions (Vollmer et al., 2019). In the first step of transpeptidation, the active site serine of PBPs makes a nucleophilic attack on the peptide bond between D-Ala-D-Ala of a pentapeptide producing a covalent serine-acyl intermediate and release of the terminal D-Ala. Next, the epsilon amino group of L-Lys of a peptide stem of an adjacent glycan chain attacks the serine-acyl bond resulting in a direct peptide crosslinking (Blumberg & Strominger, 1974; Joris et al., 1988; Tipper & Strominger, 1968). In *S. pneumoniae* the di-peptides L-Ser-L-Ala or L-Ala-L-Ala can be added to the epsilon amino group of pentapeptide lysine residues before transpeptidation. When such branched stem peptides are used in transpeptidation reactions, the crosslink will contain the L-Ser/Ala-L-Ala di-peptide between the D-Ala and L-Lys (Filipe et al., 2000; Lloyd et al., 2008). Since β-lactams structurally mimic the D-Ala-D-Ala moiety, they will bind to the active transpeptidation site of PBPs and form a stable serine-acyl complex thereby permanently blocking the enzyme function (Blumberg & Strominger, 1974; Frere et al., 1976; Spratt, 1975; Tipper & Strominger, 1965). β-lactam resistant pneumococci, however, express mutated *pbp* genes encoding so-called low-affinity PBPs that have reduced affinity for β-lactams while retaining their transpeptidase function (Hakenbeck et al., 2012). This allows pneumococci to grow when exposed to higher concentrations of β-lactam antibiotics.

Pneumococcal β-lactam resistance is found in strains simultaneously expressing low-affinity versions of the class B PBP2b and PBP2x and class A PBP1a (Hakenbeck et al., 2012). PBP2b and PBP2x are part of two distinct peptidoglycan synthesising machineries called the elongasome and the divisome, respectively (Briggs et al., 2021; Massidda et al., 2013; Pinho et al., 2013; Straume et al., 2021). The elongasome and divisome are recruited to the future division site at the cell equator (after establishment of the FtsZ ring) where they synthesise the primary peptidoglycan leading to cell elongation and septation (Briggs et al., 2021). In the elongasome PBP2b works alongside a dedicated shape, elongation, division and sporulation (SEDS) glycosyl transferase called RodA to synthesise peripheral peptidoglycan responsible for cell elongation. Similarly, PBP2x is part of the divisome where it together with the SEDS polymerase FtsW synthesises the septal peptidoglycan that separates the two daughter cells (Berg et al., 2013; Cho et al., 2016; Emami et al., 2017; Meeske et al., 2016; Perez et al., 2021; Perez et al., 2019; Sjodt et al., 2020; Tsui et al., 2014). The combined actions of the elongasome and divisome result in the ovoid shape of pneumococcal cells. Reduced elongasome activity results in chains of short cells that are separated by septal crosswalls. Inhibition of the divisome on the other hand results in elongated cells without septal crosswalls (Berg et al., 2013). While PBP2x and PBP2b are essential enzymes, a Δ*pbp1*a mutant is viable (Land & Winkler, 2011; Paik et al., 1999). Evidence suggests that PBP1a is part of the elongasome possibly involved in maturation of the primary peptidoglycan layer and repair of damaged peptidoglycan (Land et al., 2013; Land & Winkler, 2011; Paik et al., 1999; Perez et al., 2024; Straume et al., 2020; Tsui et al., 2014).

Although low affinity versions of PBP2x, PBP2b and PBP1a are the main drivers of penicillin resistance in pneumococci, recent decades of research have revealed several non-PBP mediated resistance factors in *S. pneumoniae* (Chesnel et al., 2005; Crisóstomo et al., 2006; Grebe et al., 1997; Guenzi et al., 1994; Huang et al., 2018; Kobras et al., 2023; Sauerbier et al., 2012; Schweizer et al., 2017; Smith & Klugman, 2001; Soualhine et al., 2005; Todorova et al., 2015; Tran et al., 2011). Among the most recent is Pde1, a phosphodiesterase that hydrolyse the second messenger molecule cyclic di-adenosine monophosphate (cyclic di-AMP) (Kobras et al., 2023). Mutations in the *pde1* gene resulting in a non-functional enzyme as well as a Δ*pde1* mutant have been shown to increase ampicillin resistance in pneumococci (Kobras et al., 2023). The same study also showed that *pde1* mutations were more prevalent in resistant isolates. Furthermore, homologs to Pde1 (usually named GdpP for <u>G</u>GDEF <u>d</u>omain <u>p</u>rotein containing <u>p</u>hosphodiesterase in other species) have been shown to be involved in β-lactam resistance in many Gram-positive species including *Staphylococcus aureus* (Argudín et al., 2016; Argudín et al., 2018; Banerjee et al., 2010; Corrigan et al., 2011; Griffiths & O’Neill, 2012; Jia et al., 2026; Lai et al., 2024; Poon et al., 2022; Sommer et al., 2021), *Bacillus subtilis* (Luo & Helmann, 2012) *Listeria monocytogenes* (Massa et al., 2020; Witte et al., 2013), *Streptococcus mutans* (Cheng et al., 2016) and *Lactococcus lactis* (Smith et al., 2012).

Intracellular cyclic di-AMP levels are controlled by the combined action of diadenylate cyclases and phosphodiesterases. In pneumococci, cyclic di-AMP is produced by the essential diadenylate cyclase CdaA (previously named DacA), which converts two molecules of ATP to cyclic di-AMP (Bai et al., 2013). The level of cyclic di-AMP can be reduced by the phosphodiesterases Pde1 and Pde2 (**Figure 1A**). Pde1 converts cyclic di-AMP to phosphoadenylyl adenosine (pApA), which can be further hydrolysed to two AMPs by Pde2. Pde2 can also hydrolyse cyclic di-AMP directly to produce two AMP molecules (Bai et al., 2013). Thus, β-lactam resistant pneumococci with a compromised Pde1 have reduced ability to lower the intracellular levels of cyclic di-AMP, which could accumulate and influence a wide range of signalling processes. Cyclic di-AMP has been shown to regulate potassium transport, osmotic and acidic stress, DNA repair, sporulation, genetic competence, biofilm formation and virulence, in addition to β-lactam resistance (reviewed in Zarrella and Bai (2020) and Yin et al. (2020)). The best-studied function in pneumococci is how cyclic di-AMP regulates turgor pressure via potassium uptake. High levels of cyclic di-AMP have been shown to inhibit potassium uptake resulting in reduced turgor pressure. Specifically, the TrkH potassium transporter is regulated by the <u>c</u>yclic di-<u>A</u>MP <u>b</u>inding <u>p</u>rotein CabP. In the unbound state CabP interacts with and activates TrkH, but upon binding to cyclic di-AMP, the interaction with TrkH is lost thereby leading to reduced potassium uptake (Bai et al., 2014). Similar regulation of potassium is observed in many other species, recently reviewed by Foster et al. (2024). The link between increased β-lactam resistance and mutations in *pde1* has therefore been attributed to regulation of turgor pressure, in which increased levels of cyclic di-AMP (from mutations in *pde1*) result in reduced turgor and thereby increased tolerance to peptidoglycan synthesis inhibition. In addition, some studies report that deletion of genes encoding cyclic di-AMP phosphodiesterases changes the peptidoglycan precursor pool and structure of peptidoglycan, suggesting that the function of PBPs and/or cell wall remodelling proteins are influenced (Corrigan et al., 2011; Dengler Haunreiter et al., 2023; Massa et al., 2020; Zhu et al., 2016). The precise molecular mechanism by which cyclic di-AMP influences β-lactam resistance therefore remains unsolved.

**Figure 1:**
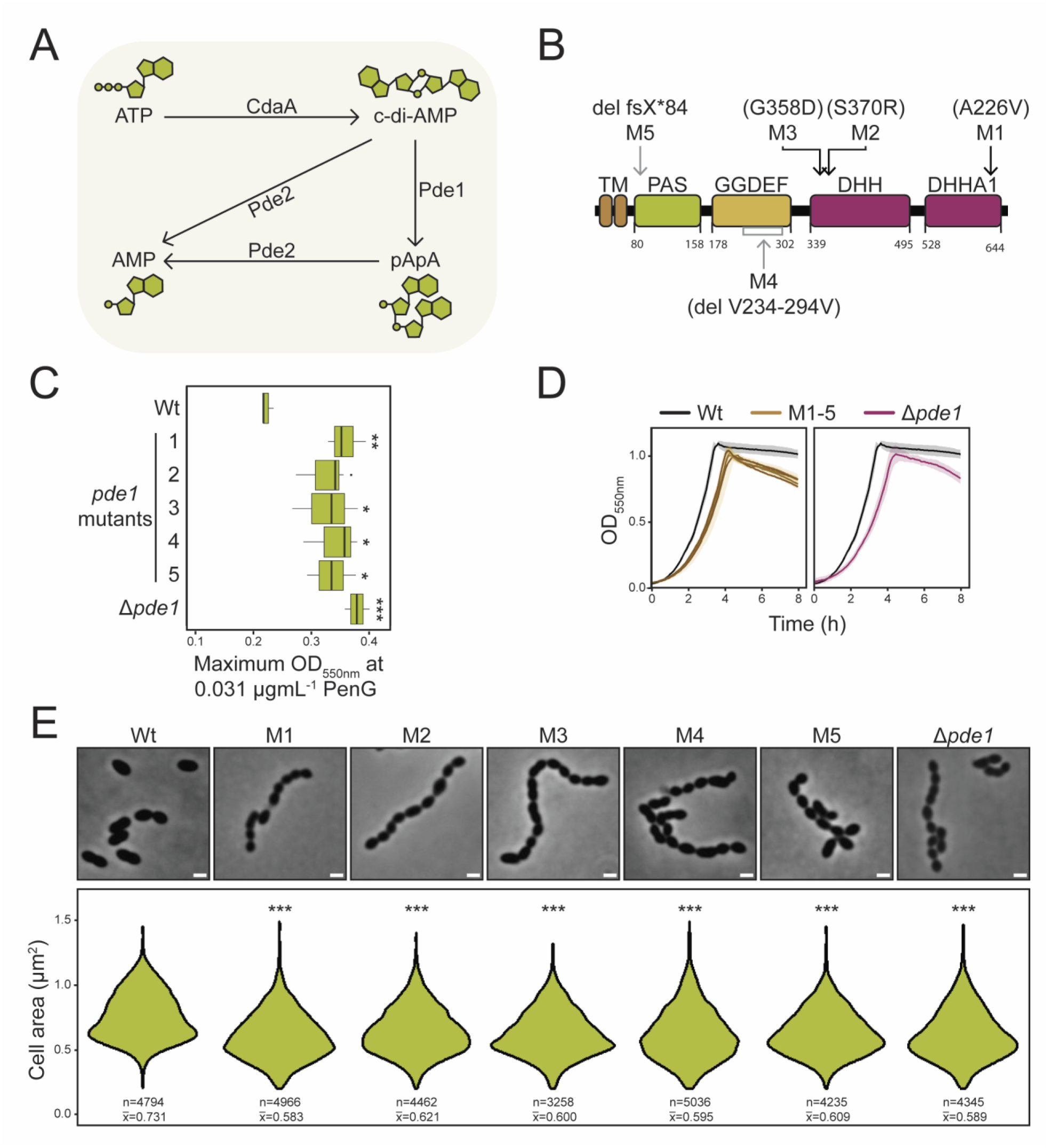
Effects of high cyclic di-AMP levels on penicillin susceptibility, growth and morphology. **(A)** A schematic overview of the cyclic di-AMP (c-di-AMP) homeostasis in *S. pneumoniae*. Cyclic di-AMP is produced from two molecules of ATP by CdaA, converted to phosphoadenylyl adenosine (pApA) by Pde1 and further hydrolysed to AMP by Pde2. Pde2 is also reported to convert cyclic di-AMP directly to AMP *in vitro*. **(B)** Illustrations of the Pde1 domain organisation and the respective protein changes identified in the *pde1* mutants (M1-5). Deletion (del), frameshift (fsX) and stop codon (*) are indicated. **(C)** Maximum growth of Wt, *pde1* mutants (M1-5) and a Δ*pde1* mutant at 0.031 µg × mL^−1^ Penicillin G (PenG) based on three biological replicates. All mutants displayed decreased penicillin susceptibility compared to Wt. **(D)** Growth measurements (OD550nm) for Wt, *pde1* mutants (M1-5) and the Δ*pde1* mutant. The data represent average and standard deviation from three biological replicates. Both the *pde1* mutants (M1-5) and the Δ*pde1* mutant displayed slower growth compared to the Wt. **(E)** The *pde1* mutants (M1-5) and the Δ*pde1* mutant displayed more chains and smaller cells (15-20% reduction) compared to the Wt. The scale bar is 1 µm, number of cells (n) and average cell size (X̅) is indicated. Microscopy was repeated three times with similar results. Statistical significance compared to Wt is indicated by *** (P≤0,001), ** (P≤0,01), * (P≤0,05) and · (P≤0.1).

In this work we present evidence linking cyclic di-AMP regulation with the function of the cell division machinery in *S. pneumoniae*. We show that low cyclic di-AMP levels impaired proper division activity in such a way that cells initiated new rounds of division before completing the previous septal cross wall. In addition, we observed that cells with elevated levels of cyclic di-AMP contained less branched stem peptides relative to linear in their cell wall, suggesting that the cell wall synthesis machineries were disturbed. In addition, we present evidence suggesting that the proteins involved in cyclic di-AMP synthesis and degradation (CdaA, CdaR and Pde1) interact and they were found to locate at the division site, indicating that cyclic di-AMP regulation could be important at this location in the cells. Our data show that the level of cyclic di-AMP is important for proper cell division in *S. pneumoniae* and that mis-regulation of this second messenger could influence β-lactam resistance.

## Results

### Loss of function mutations in *pde1* result in decreased penicillin susceptibility in *S. pneumoniae*

To identify non-*pbp* factors contributing to penicillin non-susceptibility in *S. pneumoniae*, we initially transformed the penicillin susceptible *S. pneumoniae* strain R6 (hereafter referred to as wild-type [Wt]) with a pool of DNA fragments containing PCR amplicons of the genes *ftsL*, *mraY*, *murE*, *dltA*, *ciaH*, *clpL*, *recU* and *dhfR* from the penicillin resistant *Streptococcus oralis* Uo5. It has been found that these loci are more frequently mutated in penicillin resistant isolates (Chewapreecha et al., 2014; Todorova et al., 2015), which was also the case for the Uo5 strain. Transformants with decreased penicillin sensitivity were selected using a 1.5-fold gradient of Penicillin G (PenG). After 24 hours, a few small colonies appeared on the transformant plates containing 0.008 µg/mL PenG (and none for the negative control). Five colonies (M1-5) were selected and whole genome sequenced. No mutations were found in the genes used as transforming DNA, but they had mutations in the phosphodiesterase encoding gene *pde1*. Pde1 has a DHH (Asp-His-His motif) and a DHHA1 (<u>DHH</u>-<u>a</u>ssociated domain <u>1</u>) domain, which are shown to be essential for phosphodiesterase activity (Bai et al., 2013; Corrigan et al., 2011; Rao et al., 2010). Additionally, it contains two transmembrane helices, a PAS (<u>P</u>er-<u>A</u>rnt-<u>S</u>im) domain possibly involved in a sensory mechanism and a degenerate GGDEF domain (Bai et al., 2013). Proteins with a GGDEF motif are commonly associated with synthesis or hydrolysis of cyclic di-GMP (Ryjenkov et al., 2005; Tal et al., 1998). Among the five mutations in *pde1*, three were mis-sense mutations in the DHH/DHHA1 domains (M1-3), one resulted in a partial deletion of the GGDEF domain (M4) and the last had a deletion causing a truncation of the protein (M5) (**Figure 1B**). The mutants M1-5 displayed decreased penicillin susceptibility compared to Wt (**Figure 1C**). We further constructed a Δ*pde1* mutant which displayed similar decrease in penicillin susceptibility (**Figure 1C**), indicating that mutants M1-5 express loss-of-function versions of the Pde1 enzyme. Additional to the effect on penicillin susceptibility, the *pde1*-mutants and the Δ*pde1* mutant displayed reduced growth and smaller cell sizes (15-20% reduction) compared to Wt (**Figure 1D, E**). We also constructed Δ*pde1* mutants in ten penicillin non-susceptible or resistant clinical isolates of *S. pneumoniae*, which resulted in increased penicillin resistance and reduced growth (**Figure S1**). Our results align with Kobras et al. (2023), who found mutations in *pde1* when growing cells on subinhibitory concentrations of ampicillin. For a comprehensive characterization of the mutation hotspots in the pneumococcal *pde1* and the prevalence of these mutations in β-lactam resistant clinical isolates, we refer to the work of Kobras et al. (2023).

### The influence of cyclic di-AMP levels on cell division and penicillin susceptibility

Given the link between penicillin susceptibility and *pde1*, combined with the observation that *pde1* deletion resulted in smaller cell sizes, we next aimed to unravel how different levels of cyclic di-AMP affect the pneumococcal cells. It is known that Δ*pde1* results in decreased cyclic di-AMP degradation and lack of pApA resulting in decreased cell size (Bai et al., 2013; Kobras et al., 2023). In order to monitor cyclic di-AMP levels in the different mutants, we developed a reporter assay inspired by Brogan et al. (2025) in which a cyclic di-AMP riboswitch upstream the *B. subtilis kimA* gene controls translation of a luciferase encoding gene (**Figure 2A**). Consequently, luciferase activity is reduced in cells with high cyclic di-AMP levels and increased in cells with low cyclic di-AMP levels. We also measured, cyclic di-AMP levels using a competitive ELISA assay for selected mutants (**Figure S2**). As expected, both the reporter assay and ELISA confirmed higher cyclic di-AMP levels in the Δ*pde1* mutant (**Figure 2B, C**, **Figure S2**). Likewise, overexpression of the cyclic di-AMP synthase CdaA increased cyclic di-AMP levels (**Figure 2B, C**), and similar to the Δ*pde1* mutant (**Figure 1C, E**), this resulted in smaller cells (**Figure 2D, E**) and reduced penicillin susceptibility (**Figure 2F**).

**Figure 2:**
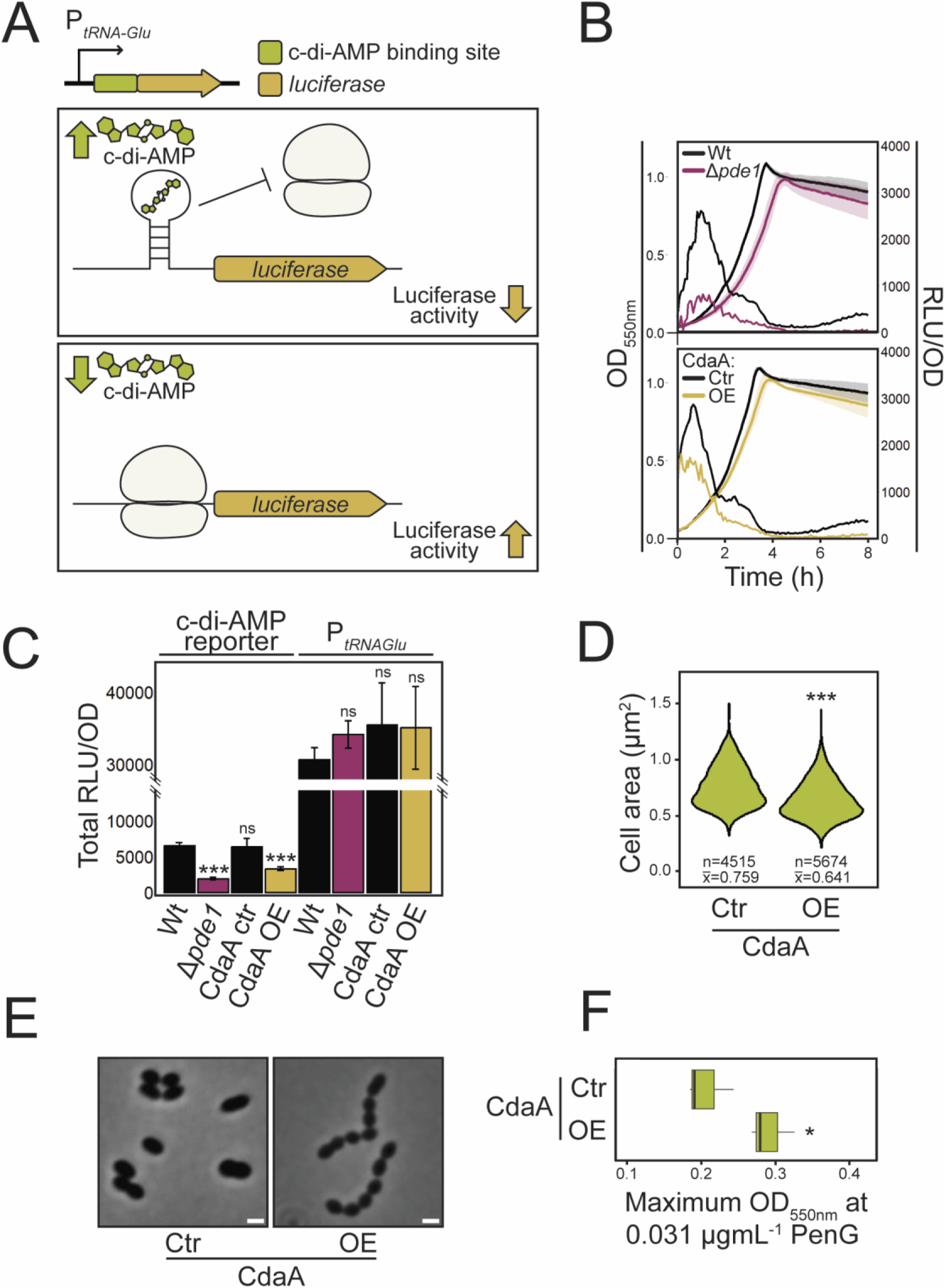
Cyclic di-AMP levels influence cell size and penicillin susceptibility. **(A)** Illustration of the cyclic di-AMP (c-di-AMP) reporter mechanism. A cyclic di-AMP riboswitch controlling mRNA translation was placed upstream a luciferase encoding gene. High concentrations of cyclic di-AMP will bind to the riboswitch upstream of the ribosomal binding site and inhibit ribosome binding, consequently reducing the amount of luciferase in the cells. Luciferase activity is therefore inversely proportional to the cyclic di-AMP levels in the cells. **(B)** Growth (OD550nm) and luciferase activity (RLU) relative to OD were measured over time in a Δ*pde1* and *cdaA* overexpression (OE) mutant. Both displayed lower luciferase signal (thereby higher cyclic di-AMP levels) than Wt and the overexpression control, respectively. The curves represent average measurements from three biological replicates, and standard deviations are indicated. **(C)** Total area under the RLU/OD curves showing a significant decrease in luciferase activity for Δ*pde1* and *cdaA* overexpression cells compared to Wt. The activity of the promoter without the riboswitch was measured as a control and were not significantly (ns) different in the mutants. **(D-E)** Cells overexpressing *cdaA* showed 12% reduced size and more chains compared to the control. Number of cells (n) and average cell size (X̅) is indicated. The microscopy was repeated three times with similar results (Scalebar = 1 µm). **(F)** Overexpression of CdaA resulted in decreased penicillin susceptibility compared to the control, based on three biological replicates. Statistical significance is indicated by *** (P≤0,001), * (P≤0,05) and ns (P>0.1).

Since increased cyclic di-AMP levels are associated with reduced cell size and decreased penicillin susceptibility in pneumococci, we wondered if lowering the cyclic di-AMP levels would have the opposite effects. Indeed, when the cyclic di-AMP levels were reduced by overexpressing *pde1*, penicillin susceptibility increased and the cells became larger (**Figure 3A, B, C**). Surprisingly, these long, enlarged cells displayed multiple incomplete septa (**Figure 3D**). Likewise, depletion of *cdaA* expression also resulted in similar morphology as *pde1* overexpression (**Figure 3D**, also reported by Jana et al. (2024)). Taken together, these data suggest a link between cyclic di-AMP levels and cell division.

**Figure 3:**
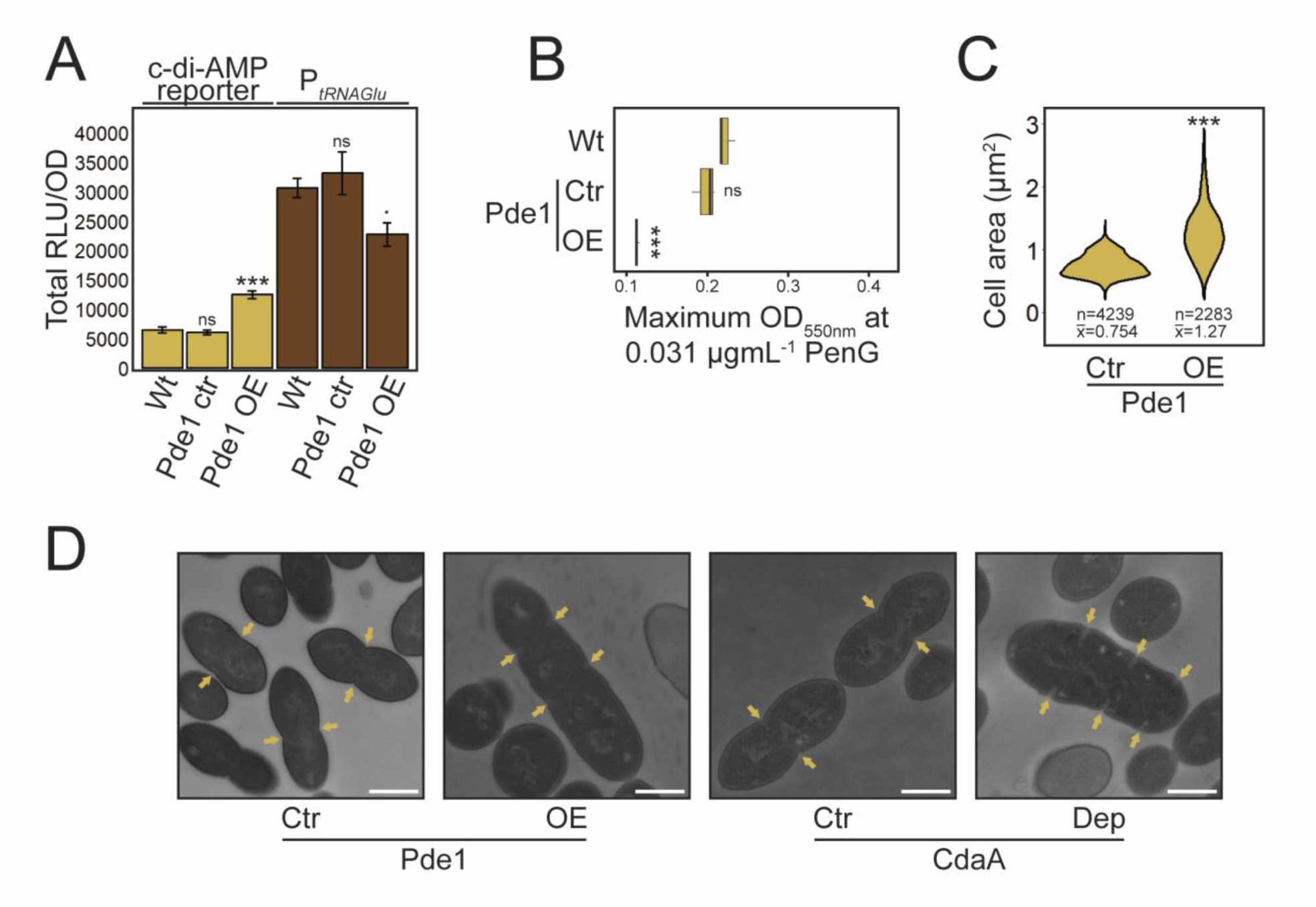
Reduced levels of cyclic di-AMP lead to elongated cells with multiple incomplete septa. **(A)** The cyclic di-AMP levels were measured using the cyclic di-AMP riboswitch reporter system (see Figure 2A). Overexpression (OE) of *pde1* showed significantly higher luciferase activity (corresponding to lower cyclic di-AMP levels) than the Wt and control (non-induced), even though the activity of the promoter (P*tRNAGlu*) was reduced (P≤0.1). Hence, the decrease in cyclic di-AMP levels might be even larger than measured with this reporter system. The results are mean and standard deviation of three biological replicates. **(B)** Overexpression of *pde1* showed higher penicillin susceptibility than the Wt and control (based on three biological replicates). Statistical significance is indicated by *** (P≤0,001), · (P≤0.1) and ns (P>0.1), respectively. **(C)** Cell area represented as violin plots showing that *pde1* overexpression resulted in an average increase in cell size of 74%. **(D)** Transmission electron microscopy of *pde1* overexpression and CdaA depleted (Dep) cells. Septa are indicated with arrows, and scale bars represent 500 nm in size.

Similar to the Δ*pde1* mutant, we expected that cells deficient of the second phosphodiesterase, Pde2, would accumulate cyclic di-AMP. Results from the competitive ELISA assay suggested that the Δ*pde2* mutant contained ∼5-fold more cyclic di-AMP compared to Wt (Δ*pde1* mutant gave an ∼11-fold increase compared to Wt) (**Figure S2**). However, when using the *in vivo* reporter assay no change in cyclic di-AMP levels was observed in Δ*pde2* cells (**Figure 4A**) and the mutant did not show reduced penicillin susceptibility as observed for Pde1 deficient cells (**Figure 4B**). Similarly, overexpression of *pde2* gave no significant changes in cyclic di-AMP levels (**Figure 4A**) and resulted in only marginally larger cells (**Figure 4C**, 27% increase, while *pde1* overexpression resulted in a 74% increased cell size). Finally, deletion of *pde2* did not further elevate the cyclic di-AMP levels of Δ*pde1* cells (**Figure 4A**, **Figure S2**). Thus, even though a Δ*pde2* mutant might have increased cyclic di-AMP levels (based on the ELISA assays), our results suggest that Pde1 is the main cyclic di-AMP hydrolysing enzyme in pneumococci. Although Pde2 might have a less prominent role in cyclic di-AMP hydrolysis, it is still important to the cells, since a Δ*pde2* mutant showed reduced growth (**Figure 4D**) and displayed abnormal cell morphology (**Figure 4E**).

**Figure 4:**
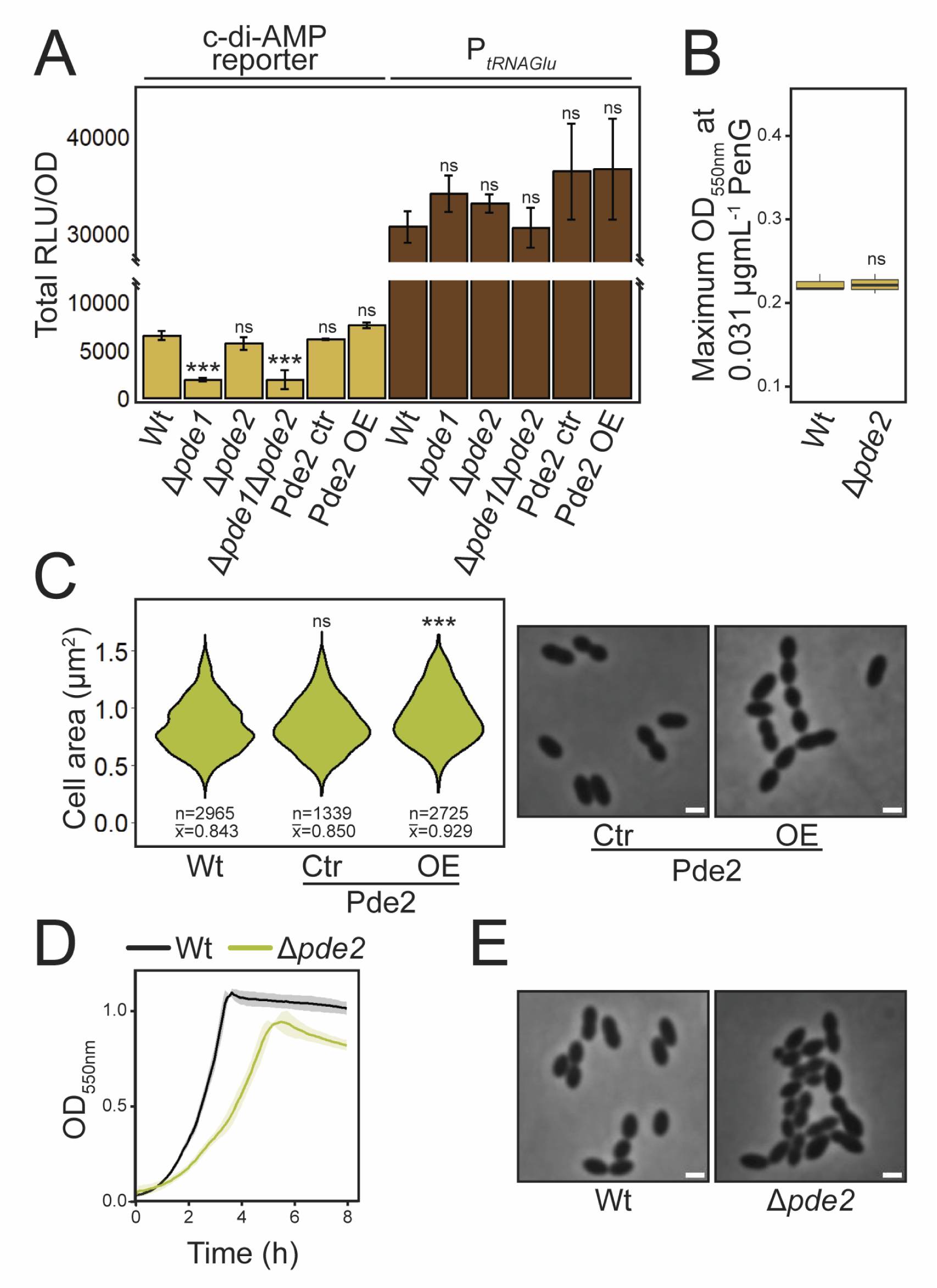
Phenotypic effects of deleting and overexpressing *pde2*. **(A)** Cyclic di-AMP levels were measured using the cyclic di-AMP riboswitch reporter system (see Figure 2A). The Δ*pde1* and double Δ*pde1*Δ*pde2* mutants had significantly lower luciferase activity (higher cyclic di-AMP level) than the Wt, while Δ*pde2* and overexpression of *pde2* displayed no significant (ns) changes compared to the Wt. The promoter activity (P*tRNAGlu*) of the mutants was not significantly changed. Data were obtained from three biological replicates. **(B)** PenG susceptibility of a Δ*pde2* mutant compared with the parental Wt. **(C)** Violin plots of cell area and phase contrast images of cells overexpressing *pde2* compared with Wt and non-induced control cells. P≤0.001 and P>0.1 are indicated by *** and ns, respectively. Scalebar = 1 µm, number of cells (n) and average cell size (X̅) is indicated. **(D)** Growth comparison of a Δ*pde2* mutant and Wt (data from three biological replicates). **(E)** Phase contrast images of a Δ*pde2* mutant and Wt. The microscopic imaging was repeated three times with similar results, scale bar = 1 µm.

### Enzymes involved in cyclic di-AMP synthesis and degradation localise to the division site

Since perturbed levels of cyclic di-AMP interfere with normal cell division, we were curious whether proteins involved in cyclic di-AMP metabolism would display distinct cellular localizations similar to typical cell division proteins. To our knowledge, the cellular localization of these proteins or their homologues has not been studied. Interestingly, using (time-lapse) fluorescence microscopy, we found that both Pde1-GFP and CdaA-GFP have a dynamic localization in the membrane with strongly enriched signals at mid-cell (**Figure 5A, B**, **Video S1, S2**). On the other hand, Pde2-GFP showed uniform intracellular location (**Figure 5B**). Using a bacterial two-hybrid system (BACTH) we tested for possible interactions between CdaA, Pde1 and Pde2. The results suggested that all three interact with each other (**Figure 5C**). Our data thus indicate that both cyclic di-AMP synthesis (CdaA) and degradation (Pde1) are enriched close to the division machinery.

**Figure 5:**
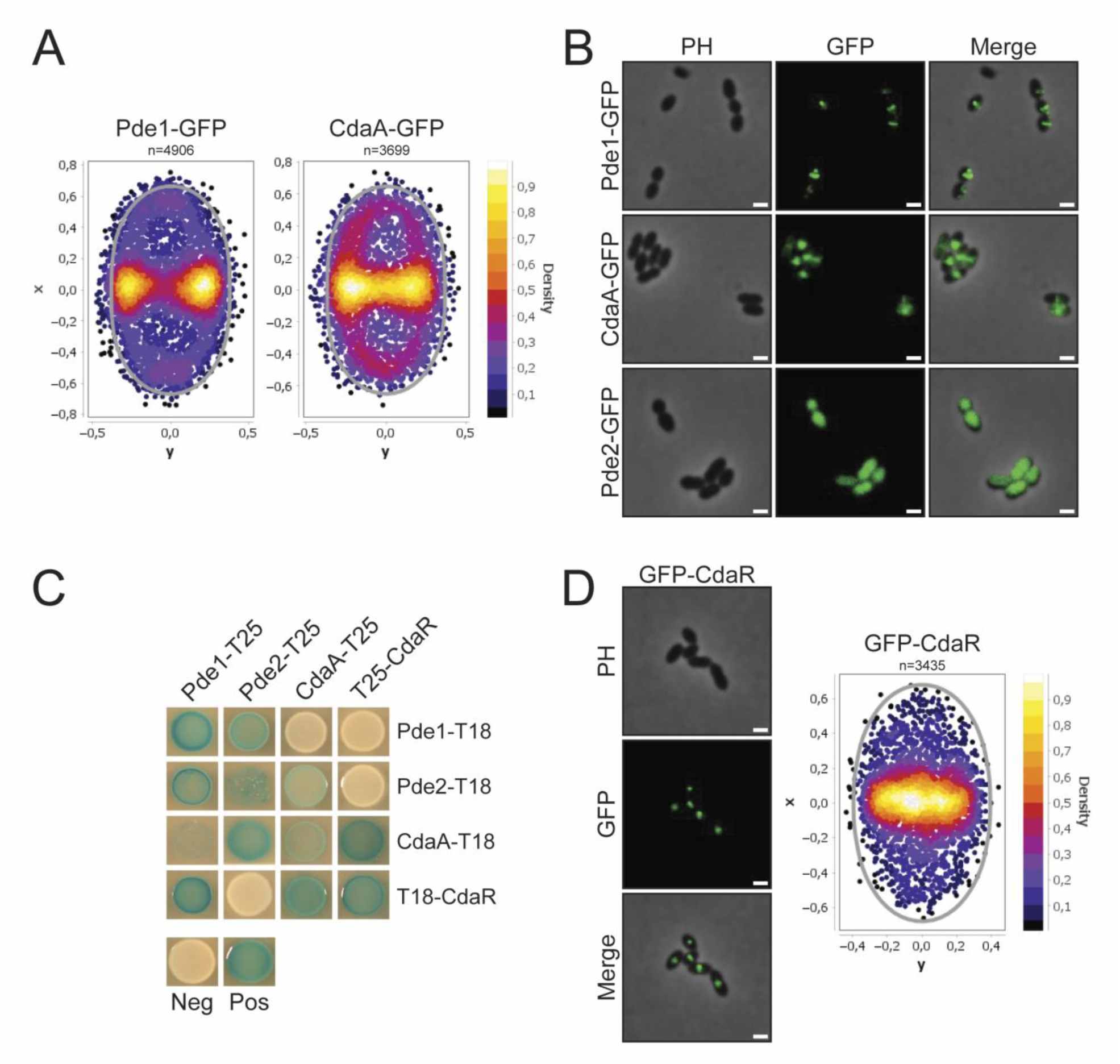
Proteins involved in cyclic di-AMP homeostasis interact and localize to mid-cell. **(A)** Density plots of the fluorescence maxima of Pde1-GFP and CdaA-GFP showed signal enrichment at mid-cell. The fluorescence signals showed a dynamic pattern as visualised in supplemental Video S1 and S2. **(B)** Fluorescence microscopy images showing localization of Pde1-GFP, CdaA-GFP and Pde2-GFP. The microscopy was repeated three times with similar results, scalebar = 1 µm. **(C)** Bacterial two hybrid system (BACTH) testing interactions between Pde1, Pde2, CdaA and CdaR. Positive interactions produce blue spots of bacteria. **(D)** Representative fluorescence microscopy image of GFP-CdaR localisation and density plot showing that the maxima fluorescence signals were focused at mid-cell.

### Deciphering the cyclic di-AMP regulatory network in *S. pneumoniae*

In several species, including *S. aureus*, *B. subtilis, L. lactis* and *L. monocytogenes*, CdaA (also called DacA or YbbP) activity has been found to be controlled by CdaR (also called YbbR) and GlmM, encoded on the *cdaA-cdaR-glmM-*operon (Brogan et al., 2025; Gundlach et al., 2015; Mehne et al., 2013; Rismondo et al., 2016; Tosi et al., 2019; Zhu et al., 2016). This genetic organization is conserved in many bacteria, including *S. pneumoniae* (Bai et al., 2013; Mehne et al., 2013). We therefore hypothesized that CdaR is involved in cyclic di-AMP regulation also in *S. pneumoniae*. First, we used BACTH analysis to test for protein-protein interactions, which suggested that CdaA and CdaR interact (**Figure 5C**), in agreement with works in other bacteria (Brogan et al., 2025; Mehne et al., 2013; Rismondo et al., 2016). Our assays also suggest that CdaR interacts with Pde1, but not Pde2 (**Figure 5C**). Next, we tested the localization of GFP-CdaR and found that it also localizes to mid-cell, but the signal was not dynamic as for Pde1-GFP and CdaA-GFP (**Figure 5D**).

Based on the results in *B. subtilis* and *L. monocytogenes,* CdaR is hypothesised to sense the state of the peptidoglycan layer and regulate the cyclic di-AMP synthesis activity of CdaA accordingly (Brogan et al., 2025; Rismondo et al., 2016). We depleted *cdaR* expression in *S. pneumoniae* and found that these cells became elongated and displayed multiple incomplete septa resembling CdaA depleted cells, indicating that CdaA is activated by CdaR (**Figure 6A, B**). Interestingly, after prolonged depletion of *cdaR* expression, the phenotype shifted from the elongated shape with multiple septa to cells with spherical shapes in chains. Together, this indicates that CdaR have an intricate regulatory effect on cell division. To make sure that the shifting phenotype was not caused by compensatory mutations, we diluted the culture of severally depleted spherical cells and added the inducer to reestablish expression of *cdaR*. These cells regained the Wt phenotype (**Figure 6B**), supporting that both phenotypes were caused by CdaR depletion and not secondary mutations. Additionally, we also spotted dilutions of CdaR depleted cells on agar plates with and without inducer and found that they only grew when *cdaR* expression was induced (**Figure 6C**). Interestingly, however, a few small colonies appeared on the plate without inducer after prolonged depletion, potentially representing suppressor mutations allowing the cells to survive in the absence of CdaR. Whole genome sequencing of four potential *cdaR* suppressor mutants (S1-4) identified that S1 had a mutation in *cdaA*, and S2-4 had mutations in *pde1* (**Table S1**). In line with this, the mutants displayed reduced cell size (**Figure S3**), another indication of elevated cyclic di-AMP levels. This shows that *S. pneumoniae* can survive without CdaR if they find other ways to regulate cyclic di-AMP homeostasis.

**Figure 6:**
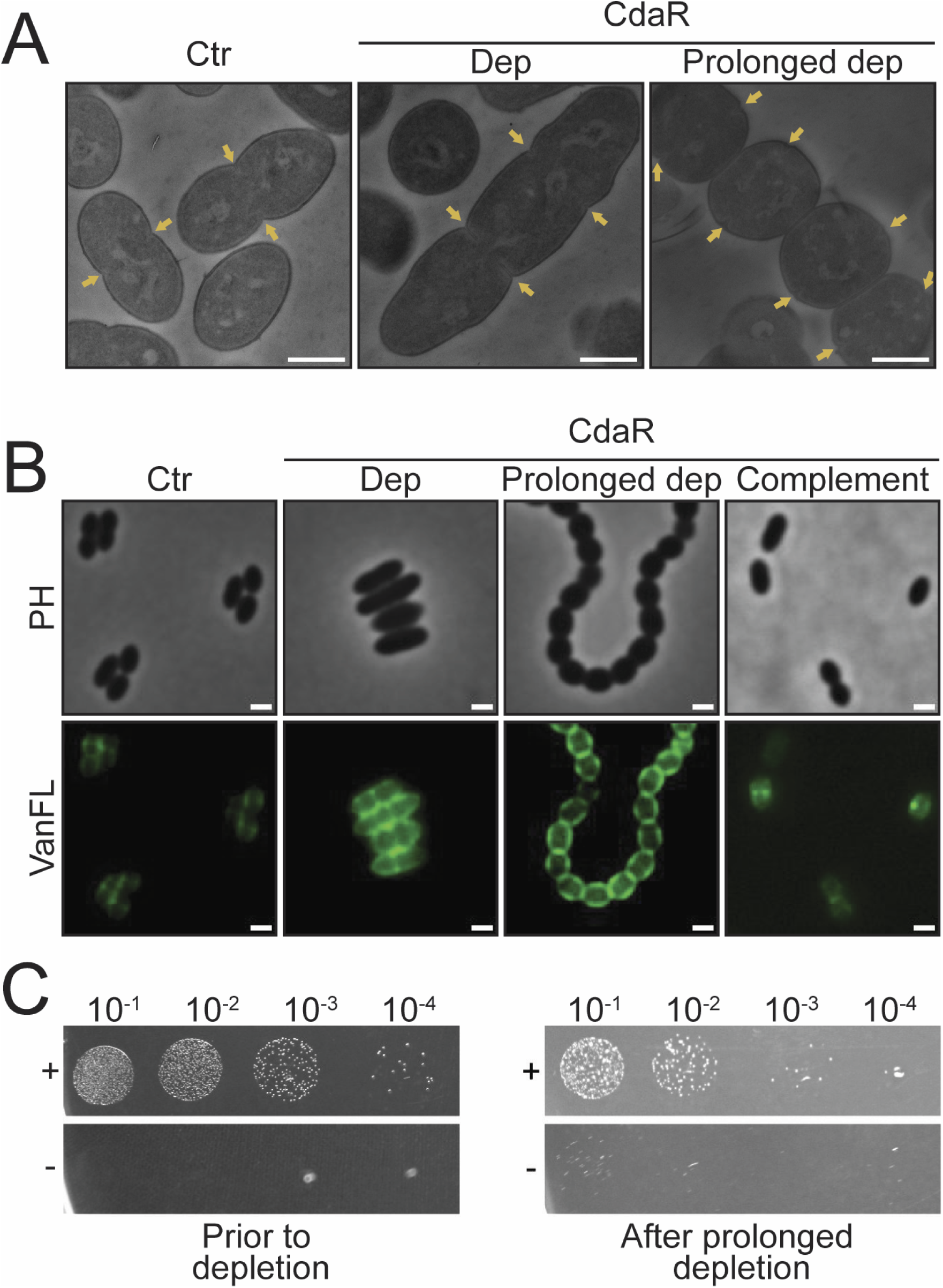
Sequential morphology changes upon CdaR depletion. Transmission electron microscopy **(A)** and fluorescence microscopy (VanFL) with corresponding phase contrast images **(B)** of *S. pneumoniae* depleted (Dep) and prolonged depleted of CdaR. Depleted cells (grown 2 hours without ComS inducer) displayed large, elongated cells with multiple incomplete septa (indicated with arrows), while after prolonged depletion (grown 5.5 hours without ComS inducer) the morphology shifted to a phenotype of spherical cells growing in long chains. The Wt phenotype was restored when the cells of prolonged depletion were grown in medium containing inducer (complementation). Scale bars are 500 nm for electron micrographs and 1 µm for fluorescence images. **(C)** A 10-fold dilution series of the CdaR depletion strain spotted on agar plates with (+) and without (-) inducer (left panel). After prolonged depletion, the culture was also diluted and spotted onto agar plates (right panel).

### Cyclic di-AMP levels influence the cell wall composition and PBP localization

It has been observed in several species (*S. aureus*, *L. lactis*, *B. subtilis* and *L. monocytogenes*) that the cyclic di-AMP levels can influence cell wall synthesis (Corrigan et al., 2011; Luo & Helmann, 2012; Massa et al., 2020; Zhu et al., 2016). We therefore analysed the cell wall stem peptide composition of pneumococcal Δ*pde1* and Δ*pde2* mutants. The Δ*pde1* mutant displayed a cell wall consisting of more linear stem peptides relative to branched compared to the Wt, while the Δ*pde2* mutant was more similar to Wt (**Figure 7A, B**). A recent study by Jia et al. (2026) reported that high cytoplasmic levels of cyclic di-AMP somehow caused PBP mis-localization in *S. aureus*. We tested whether this was also the case for pneumococci lacking *pde1* by labelling PBPs with the fluorescent penicillin Bocillin FL. The number of cells with Bocillin FL foci increased by 400% in the Δ*pde1* mutant compared to Wt (**Figure 7C**), suggesting that this effect observed in *S. aureus* (Jia et al., 2026) is conserved among different bacterial species.

**Figure 7:**
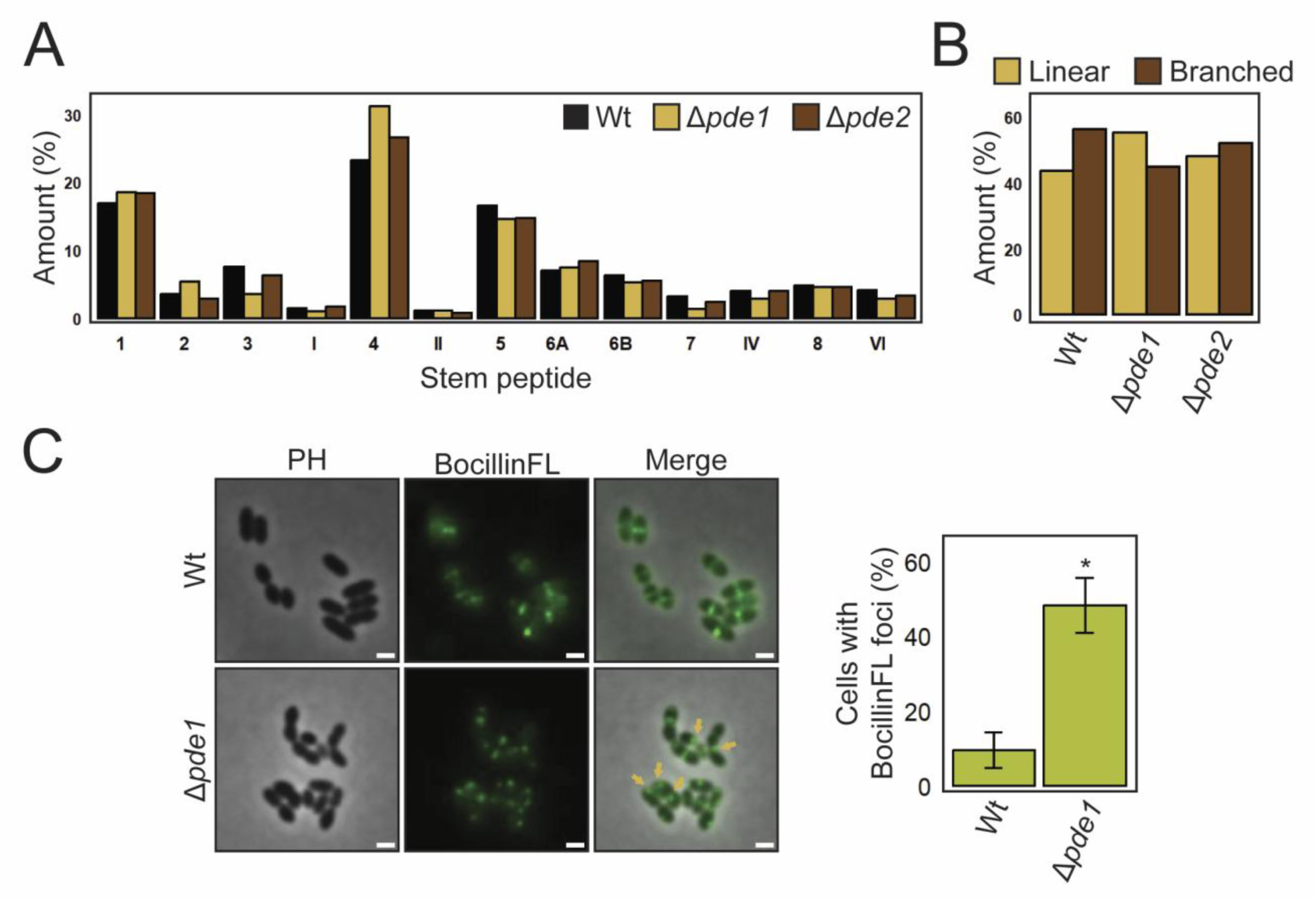
Stem peptide composition and PBP localization are affected by cyclic di-AMP levels. **(A)** Comparison of relative percentage of each stem peptide in the peptidoglycan of Wt-, Δ*pde1*-, and Δ*pde2*-cells. Linear monomers (1 and 2), branched monomers (3, I, II), linear dimer (4) and branched dimers (5, 6A, 6B, 7, IV, 8, VI). Overview of stem peptide structures is shown in **Figure S6**. **(B)** Comparison of linear versus branched stem peptides in the peptidoglycan of Wt-, Δ*pde1*-, and Δ*pde2*-cells. **(C)** Fluorescent signal of BocillinFL labelled PBPs in Wt and Δ*pde1* cells. Whithe Wt mostly have fluorescent signal across the mid-cell, the Δ*pde1* mutant displayed increased amounts of foci outside of mid-cell (indicated by arrows). The number of misplaced foci per 300 cells were manually counted. The bars and error bars represent mean and standard deviation calculated from two biological replicates, respectively * (P≤0,05). Scalebars are 1 µm.

## Discussion

Inactivation of phosphodiesterase enzymes has been found to increase β-lactam resistance in many Gram-positives (Argudín et al., 2016; Argudín et al., 2018; Banerjee et al., 2010; Cheng et al., 2016; Corrigan et al., 2011; Griffiths & O’Neill, 2012; Jia et al., 2026; Lai et al., 2024; Luo & Helmann, 2012; Massa et al., 2020; Poon et al., 2022; Smith et al., 2012; Sommer et al., 2021; Witte et al., 2013), including a recent study in which this phenotype was observed in *S. pneumoniae* (Kobras et al., 2023). It has been suggested that high levels of cyclic di-AMP reduce β-lactam susceptibility via reduced potassium import, which in turn results in reduced cell size and reduction in turgor pressure, making the cells more prone to resist damage to the cell wall for example implicated by cell wall targeting antibiotics (Foster et al., 2024). In the present study we have confirmed that penicillin susceptibility in *S. pneumoniae* depends on the intracellular levels of cyclic di-AMP, i.e. elevated levels decreased penicillin susceptibility, while lower levels had the opposite effect. We further studied the cellular effects of cyclic di-AMP levels and the enzymes involved in cyclic di-AMP synthesis and turnover and found that perturbations of the cyclic di-AMP levels influenced pneumococcal cell morphology in two ways: (i) higher levels of cyclic di-AMP (deletion of *pde1* or overexpression of *cdaA*) were associated with reduced cell size (**Figure 1E**, **Figure 2D**, **E**) and (ii) lower cyclic di-AMP levels (depletion of *cdaA* or overexpression of *pde1*) resulted in elongated cells with multiple incomplete septa (**Figure 3D**). This shows that high and low levels of cyclic di-AMP have opposite effects on cell morphology suggesting that the cyclic di-AMP turnover could be involved in regulation of cell division.

The elongation and incomplete septa of cells with low cyclic di-AMP levels indicate a compromised divisome. However, the morphology of PBP2x (the main transpeptidase in the divisome) depleted cells differ from cyclic di-AMP depleted cells. First, PBP2x depleted cells are frequently associated with a characteristic lemon-shape and fewer elongated cells as observed by phase contrast microscopy (**Figure S4A**) and previously shown by Berg et al. (2013). Secondly, while cells overexpressing *pde1* or depleted of CdaA or CdaR initiate septal cross wall formation (resulting in cells with multiple incomplete septa), PBP2x depleted cells are completely inhibited from starting to synthesise a new septal cross wall (multiple incomplete septa are not observed, **Figure S4B**). Together, this could indicate that low cyclic di-AMP levels slow down the divisome machinery rather than fully inhibiting its function. Interestingly, it has been reported that deletion of *pde1* could rescue the phenotype of a *pbp2x* knockdown mutant, supporting a functional link between these two genes (Jana et al., 2024). The authors attributed this effect to cyclic di-AMP mediated inhibition of potassium import resulting in reduced turgor. It has also been proposed by Commichau et al. (2018) that changes in turgor pressure indirectly influence cell division and cell wall synthesis, which in turn influence β-lactam susceptibility. Whether cyclic di-AMP can regulate cell division proteins directly or if the cell division machinery is compromised by aberrant turgor pressure is not clear.

In addition to morphological changes, we found that the ratio of linear to branched stem peptides in the cell wall of Δ*pde1* cells increased (**Figure 7A, B**) and PBPs mis-localised (**Figure 7C**) showing that peptidoglycan synthesis machineries were affected by elevated cyclic di-AMP levels. These data are in line with studies reporting that perturbed cyclic di-AMP levels influence peptidoglycan synthesis. For example, deletion of *gdpP* (*pde1* homolog) in *S. aureus* has been shown to increase the amounts of peptide crosslinks in the cell wall (Corrigan et al., 2011) and PBPs were found to mis-localise (Jia et al., 2026). It is, however, important to mention that in the results by Jia et al. and a study by Lai et al. (2024), an increase in peptidoglycan crosslinks was not observed in *S. aureus* Δ*gdpP* cells. Moreover, deletions of *pdeA* and *pgpH* (encoding cyclic di-AMP phosphodiesterases) in *L. monocytogenes* were found to reduce the cell wall thickness and concentration of peptidoglycan precursors (Massa et al., 2020). It was therefore intriguing to observe that the proteins involved in cyclic di-AMP synthesis and degradation (CdaA, Pde1 as well as the putative CdaA regulator CdaR) localized at the division site in *S. pneumoniae* (**Figure 5A**, **B**, **D**), suggesting that cyclic di-AMP signalling is particularly important at this location. Whether the potassium transport system TrkH-CabP localizes to the division zone is to the best of our knowledge not known. This was not explored in the present work, but should future studies show that TrkH and/or CabP do not localize to the division site, it would strengthen the hypothesis that strict control of cyclic di-AMP production at this site could involve regulation of cell division proteins. In fact, it has been proposed that local pools of cyclic di-AMP could direct the signal to target proteins (Bai et al., 2013). However, Foster et al. (2024) argued against this: Since the diffusion rate of cyclic di-AMP from one pole to the other is calculated to be less than 100 milliseconds for a cell of 3 µm in length, localization-dependent signalling is not likely. Thus, whether there is any localization-specific cyclic di-AMP signalling involved in regulation of potassium transport and/or cell division remains to be revealed.

We also studied the role of CdaR in *S. pneumoniae*. CdaR is a membrane protein with extracellular domains, including an intrinsically disordered region (IDR) which in *B. subtilis* has been shown to somehow sense the status of the peptidoglycan and thereby activates CdaA to produce cyclic di-AMP in response to external signals (Brogan et al., 2025). Depletion of CdaR showed an interesting dual phenotype in *S. pneumoniae*. Initially, when cells were slightly depleted of CdaR, the growth rate was reduced, and cells were enlarged with multiple incomplete septa resembling CdaA depleted cells (**Figure 6A, B**). However, when CdaR was severely depleted, the cells shifted to a chaining and spherical shape (**Figure 6A, B**). This dual CdaR-dependent phenotype is different from what was observed upon CdaA depletion and could indicate that CdaR also have other functions in the cells beyond controlling CdaA. We also observed that overexpression of CdaR only marginally reduced the average cell size (4%) (**Figure S5A**) whereas penicillin susceptibility was not affected (**Figure S5B**). This contrasts with the effects of high CdaA levels, indicating that CdaA (or the signal sensed by CdaR) could be the limiting factor in this signalling process and that more CdaR in the membrane does not lead to higher activity of CdaA.

It is also worth noting that under the experimental conditions used in the present work the second pneumococcal phosphodiesterase Pde2 seems to play a less prominent role in modulating cyclic di-AMP levels and cyclic di-AMP-associated phenotypes. The Δ*pde2* mutant displayed reduced growth similar to the Δ*pde1* mutant; however it resembled Wt with respect to penicillin susceptibility and stem peptide composition (**Figure 4B**, **7A, B**). Furthermore, overexpression of Pde2 did not decrease the cyclic-di-AMP levels and only marginally increased the cell size compared to overexpression of Pde1 (**Figure 4A, C**). Bai et al. (2013) demonstrated that Pde2 degrades cyclic di-AMP *in vitro* (*V*_max_ = 48.92 nmol mg^−1^ min^−1^) but that it likely favours pApA as substrate (*V*_max_ = 333 500 nmol mg^−1^ min^−1^), while Pde1 showed higher degradation rate of cyclic di-AMP (*V*_max_ = 101.6 nmol mg^−1^ min^−1^). These results support our data that Pde2 might not have efficient cyclic di-AMP degradation activity *in vivo*.

Whether changes in turgor pressure alone could induce the phenotypic changes observed in *S. pneumoniae* in the present study or if cyclic di-AMP is more directly involved in regulation of cell division and thereby affecting β-lactam susceptibility is still not clear. Here we show that the enzymes involved in cyclic di-AMP production and degradation interact at the site of cell division in *S. pneumoniae* and that elevated and reduced cyclic di-AMP levels have counteracting effects on cell morphology. The former produced smaller cells, while the latter resulted in elongated cells with incomplete septa suggesting that the effect of cyclic di-AMP levels on β-lactam susceptibility may involve modulations of the cell division machinery. However more studies are needed to understand the underlying mechanism of how alterations in cyclic di-AMP levels influence cell division and β-lactam susceptibility.

## Materials and Methods

### Cultivation and transformation of bacteria

Bacterial strains and plasmids used in this work are listed in supplementary **Table S2**. *S. pneumoniae* was cultured at 37°C in C-medium supplemented with yeast extract (Lacks & Hotchkiss, 1960) in closed tubes or on Todd Hewitt (TH) (Becton Dickinson) agar plates in anaerobic chambers generated by including Oxid AnaeroGen (Thermo Fisher Scientific) sachets. Transformation of *S. pneumoniae* was performed by natural transformation. Exponentially growing cultures were diluted to OD_550nm_ = 0.05 - 0.1 and incubated at 37°C for 15 minutes before competence was induced by adding a final concentration of 250 ng × mL^−1^ competence-stimulating peptide 1 (CSP-1; H_2_N-EMRLSKFFRDFILQRKK-COOH). Transforming DNA (final concentration of 100 – 200 ng × mL^−1^) was added together with CSP-1 and the cell culture was incubated at 37°C for 2 hours before plating on selective medium. For clinical isolates, the incubation time prior to competence induction was prolonged until the culture had doubled the OD_550nm_, and the incubation period post competence induction was 2.5 hours. *Escherichia coli* cells were grown aerobically at 37 °C in Lysogeny broth (LB) with shaking or on LB agar plates unless stated otherwise. For transformation of *E. coli*, chemically competent cells were prepared through calcium chloride treatment and transformation by heat shock at 42 °C for 30-40 seconds according to standard protocols. When necessary, antibiotics were added to the growth medium using the following final concentrations for *S. pneumoniae*: kanamycin (400 µg × mL^−1^), streptomycin (200 µg × mL^−1^), spectinomycin (100 µg × mL^−1^) and penicillin G (concentrations indicated when appropriate), and for *E. coli*: kanamycin (50 µg × mL^−1^) and ampicillin (100 µg × mL^−1^).

### Construction of mutants

Oligonucleotides used for PCR are listed in **Table S3**. *S. pneumoniae* was transformed with PCR amplicons containing either an antibiotic selection marker gene, a gene or sequence of interest, or a combination of both which was flanked with ∼1000 bp homologous regions upstream and downstream of the target site in the genome. Overlap extension PCR was used to construct the amplicons as previously described by Johnsborg et al. (2008) using the principle by Higuchi et al. (1988). Gene knockouts and replacements were created by using the Janus cassette (Sung et al., 2001) and the luciferase reporter system was introduced in the genome of mutants by using a downstream spectinomycin resistance cassette for selection. Detailed information of the constructs is given below. Gene knockout mutants were screened using PCR while gene replacements, and sequence insertions were confirmed using Sanger sequencing.

### Δ*pde1* and Δ*pde2* mutants

A Δ*pde1*::Janus cassette was constructed with a constitutive P1 promoter (Johnsborg & Håvarstein, 2009) at the 3’-end of *pde1* to avoid polar effects on the overlapping downstream *rpII* gene. Additionally, the last 78 bp of the 3’-end of *pde1* was left undisturbed to keep the ribosomal binding site (RSB) of *rpII*. When the Janus cassette was removed from the *pde1* locus using a Δ*pde1*::DEL construct the resulting Δ*pde1* mutant contained the P*_pde1_* promoter followed by the P1 promoter used at the end of the Janus construct and the last 78 bp sequence of *pde1*. A *pde2* knockout mutant was created by replacing the complete *pde2* ORF with the Janus cassette.

### *pde1*, *pde2*, *cdaA* and *cdaR* overexpression strains

The *pde1*, *pde2*, *cdaA* and *cdaR* genes were amplified using the RH425 (R6 derivate) genome as template before they were fused to the ∼1000 sequences flanking the janus cassette in the genome of strain SPH131, which contains the ComRS gene overexpression/depletion system (Berg et al., 2011). The ∼1000 bp upstream region contains the *cpsO* gene followed by the P*_comX_* inducible promoter and the downstream sequence contains the *cspN* gene. The resulting constructs were used to replace janus in SPH131 resulting in strains in which *pde1*, *pde2*, *cdaA* or *cdaR* were placed behind the P*_comX_* promoter.

### CdaA and CdaR depletion strains

DNA constructs to produce CdaA and CdaR depletion strains were made as described for the overexpression strains except for using a modified P*_comX_* promoter (P *^c^*), which was amplified using genomic DNA from strain SPH155 as template. The P *^c^* promoter contains a point mutation (5′-T<u>G</u>GAGGT-3′ to 5′-T<u>C</u>GAGGT-3′) in the RBS (Berg et al., 2013). Then the native *cdaA* and *cdaR* copies were deleted using the Janus cassette. To preserve integrity of operons, the Δ*cdaA* construct replaced the sequence encoding the transmembrane domains and the DAC domain (aa 25-244 of CdaA) by keeping the first 72 bp of the 5’-end of *cdaA* and the last 126 bp of the 3’-end. The same was done for *cdaR* by keeping the first 72 bp of the 5’-end of *cdaR* and the last 48 bp at the 3’-end.

### GFP fusion of Pde1, Pde2, CdaA and CdaR

GFP fusions were constructed in two different ways. For Pde1 and Pde2, the native gene was replaced by a *pde1-gfp* or *pde2-gfp* sequence. Additionally, 17 bp of the *pde1* 3’-end were added behind the *pde1-gfp* sequence to keep the RBS and 5’-end of *rpII*. While CdaA and CdaR GFP fusions were made by placing the construct under control of the P*_comX_* promoter (ComRS system). The GFP encoding sequence and the respective genes (except *pde1*) were connected by a sequence encoding a flexible linker (5’-GGATCTGGTGGAGAAGCTGCAGCTAAAGCTGGA-3’). The GFP was fused to the C-terminal end of Pde1, Pde2 and CdaA and the N-terminal end of CdaR.

### Riboswitch reporter strains

An *in vivo* cyclic di-AMP reporter was constructed using the riboswitch sequence upstream of *kimA* in *B. subtilis* (Nelson et al., 2013). We chose to place the reporter in the *zip* locus by replacing the pseudogene *spr1750* which has been proposed to be a neutral locus (Keller et al., 2019). The reporter construct consists of a strong constitutive promoter coupled to the cyclic di-AMP binding site and attenuator sequence, an RBS for translation of a luciferase encoding gene (*luc*), a second RBS followed by *gfp* encoding a monomeric sfGFP, a spectinomycin selection marker (*aad9*) and two terminator sequences (**Figure S7**, **Text S1**). The 61 bp sequence upstream the *tRNA-Glu* gene in *S. pneumoniae* was used as a strong promoter according to Shainheit et al. (2014). To make sure that the whole cyclic di-AMP binding site and attenuator sequence were included, 340 bp upstream of *kimA* in *B. subtilis* was amplified and fused to a *luc-gfp-aad9* amplicon. The *rrnB* T1 and T7Te terminators in pPEPY was added at the 3’-end to ensure termination of transcription. pPEPY was kindly given by Professor Jan-Willem Veening (Addgene plasmid # 122633; http://n2t.net/addgene:122633; RRID:Addgene_122633). Since the construct contains a translational repressor riboswitch upstream *luc*, cyclic di-AMP levels are directly correlated to differences between gene expression controlled by P*_tRNA-Glu_* and repression of *luc* translation (reduced luciferase activity). We initially planned to use the sfGFP signal as a control for stabile transcription from P*_tRNA-Glu_* since a separate RBS was placed upstream *gfp* to ensure translation independent of cyclic di-AMP levels. However, the fluorescent signal was not sufficient for this purpose. Therefore, we used the same reporter construct without the riboswitch sequence (P*_tRNA-Glu_* controlling the *luc*-*gfp* expression) to measure the activity of P*_tRNA-Glu_* (luciferase activity) as a control that the promoter activity was similar in different genetic backgrounds.

### Cyclic di-AMP reporter assay

Exponentially growing cultures were diluted in C-medium supplemented with 180 µg × mL^−1^ D-luciferin (Invitrogen) to an OD_550nm_ of 0.05 before 300 µL of the diluted culture was added to a white 96-well plate with clear flat bottom. When appropriate, ComS inducer (final concentration of 2 µM) was added to the growth medium. Both OD_550nm_ and luminescence (460 nm) were detected every 5 minutes for 20 hours using a Hidex Sense microplate reader. Relative luminescence units (RLU) were divided by OD_550nm_ values to normalise for cell density. Area under the curves between 0 and 12 hours was calculated in the Rstudio software (Posit, 2026) Version 2026.01.1 for each time series using the trapezoidal rule using the packages dplyr (Wickham et al., 2023) Version 1.1.4 and tidyr (Wickham et al., 2024) Version 1.3.1.

### Cyclic di-AMP measurements (ELISA)

Exponentially growing cultures with OD_550nm_ just above 0.5 were back-diluted to an exact volume of 20 mL at OD_550nm_ = 0.5 and cells were harvested by centrifugation for 10 minutes at 4000 *g* and resuspended in 1.2 mL C-medium. From this 1 mL culture was centrifuged at 20 000 *g* for 1 minute and the cell pellet was resuspended in 500 µL B-PERᵀᴹ Bacterial Protein Extraction Reagent (Thermo Fisher Scientific) and rotated for 30 minutes at room temperature. The lysate was stored at −20 °C until use. Each mutant was sampled three times on separate days (biological replicates) and the lysates were tested in three dilutions (with two technical replicates for each dilution). The cyclic di-AMP levels were quantified using the Cyclic di-AMP ELISA Kit from Cayman Chemical (item No. 501960) according to instructions from the manufacturer.

### Isolation and sequencing of genomic DNA

Bacteria grown to OD_550nm_ = 0.4 - 0.5 were harvested by centrifugation at 4000 *g* for 5 minutes. Genomic DNA was isolated by using NucleoBond AXG 100 columns (MACHEREY-NAGEL) and NucleoBond Buffer Set III (MACHEREY-NAGEL) as described by the manufacturer. Illumina sequencing was performed by Biomarker Technologies (BMK) and the Geneious Prime 2025.1.1 (https://www.geneious.com) software was used to process and map the reads to reference genomes for identification of mutations.

### Growth experiments and MIC determination

Exponentially growing cultures were diluted to OD_550nm_ = 0.05 in C-medium and volumes of 300 µL were transferred to a 96-well plate and OD_550nm_ was measured every 5 minutes for 16-20 hours in a Hidex Sense microplate reader at 37 °C. The plate was shaken at 300 rpm for 5 seconds prior to every measurement. MIC determination was performed by using broth microdilutions of PenG in a 96-well plate. Culture volumes of 260 µL at OD_550nm_ = 0.05 was added to 40 µL penicillin dilution series in a 96-well plate and growth was measured continuously as described. The MIC value was determined as the concentration of antibiotic that inhibited ≥50% of the average bacterial growth (maximum OD_550nm_ reached without PenG) using three biological replicates.

### Overexpression and depletion of proteins

Both overexpression and depletion of gene expression were performed using the previously described ComRS system for gene depletion in *S. pneumoniae* (Berg et al., 2011). Depletion experiments were conducted by diluting exponentially growing cultures (OD_550nm_ = 0.3 – 0.4) to an OD_550nm_ = 0.05 in C-medium containing a final concentration of 0.2 µM ComS (0.1 µM for CdaR depletion) followed by incubation at 37°C for 1.5 hours. The cells were harvested by centrifugation at 4000 *g* for 5 minutes and washed three times with C-medium to remove excess ComS. The cells were diluted to an OD_550nm_ of 0.05 in C-medium and two-fold dilution series of the culture were made in 100 µl volumes in a 96-well plate. To one 100 µL culture dilution series a volume of 200 µL of C-medium without (depletion) ComS were added to give a final volume of 300 µl in each well. As control of non-depleted cells, a parallel culture dilution series was added 200 µL C-medium containing 0.3 µM ComS (or 0.15 µM for CdaR depletion). Growth at 37°C was measured every 5 minutes for 20 hours using a Hidex Sense microplate reader. Gene overexpression was performed by including a final concentration of 2 µM ComS in the growth medium of mutants where the native gene was still intact, and the gene of interest was under control of the P*_comX_* promoter.

### Microscopy

For morphological and protein localisation experiments, phase contrast and epifluorescence microscopy were performed directly on growing cultures at OD_550nm_ = 0.3 – 0.4. Cell wall staining was done by adding a final concentration of 0.8 μg × mL^−1^ BODIPY™ FL Vancomycin (Thermo Fisher Scientific) prior to imaging. Staining of PBPs was performed by washing cells collected from 1 mL culture with 1 mL C medium prior to labelling with a final concentration of 10 µM BocillinFL for 30 minutes at 37°C in 1 mL C-medium. After incubation, the cells were harvested at 5000 *g*, washed once in 1 mL 1x PBS and resuspended in 100 µL 1x PBS prior to imaging using a Leica DMi8 microscope and a K8 Scientific CMOS camera. The images were prepared and analysed using the ImageJ software and the MicrobeJ plug-in (Ducret et al., 2016).

For transmission electron microscopy (TEM) cell from a 50 mL culture at OD_550nm_ = 0.3 were harvested at 5000 *g* for 10 minutes and the pellet was resuspended in fixative (1.25% glutaraldehyde and 2% paraformaldehyde in 0.1 M cacodylate buffer, pH 7.4) and incubated at room temperature for one hour followed by 4 °C overnight. The fixed cells were harvested by centrifugation at 8 000 *g* for 5 minutes and washed three times with 1 mL 0.1 M cacodylate buffer. The cell pellet was then embedded in 3% low-melt agarose (dissolved in MilliQ-H_2_O) and left to solidify at room temperature for 5 minutes. The pellet was submerged in 0.1 M cacodylate buffer and stored at 4 °C until further preparation of the samples. Next. the cacodylate buffer was replaced with 1 mL 1-2% Osmium (OsO_4_) Ferricyanide in 0.1M PHEM buffer and incubated for 60 minutes. The samples were washed 3 times for 10 minutes with 1 mL MilliQ-H_2_O before staining with 4% Uranyl acetate (dissolved in MilliQ-H_2_O) for 30 minutes. The samples were gradually dehydrated by 10-minutes incubations in 70, 90 and 100% ethanol before embedding in Epon epoxy resin and polymerization at 60 °C for 24 hours. The samples were sectioned, and a JEOL JEM-1230 electron microscope was used for imaging.

### Bacterial two-hybrid assay

The BACTH two-hybrid system was used to test protein interactions (Karimova et al., 1998). Genes of interest were cloned in frame with a T18 or T25 encoding sequence (using pKT25, pKNT25, pUT18 or pUT18C) to produce fusion proteins having the T18 or T25 domain either at the N-terminal or C-terminal end. The resulting plasmids (**Table S2**) were transformed into *E. coli* XL-Blue cells, and the cloned sequences were verified by Sanger sequencing. One plasmid expressing a T18 fusion and one plasmid expressing a T25 fusion were co-transformed into *E. coli* BTH101 (*cya*^−^) cells. Five colonies were grown in 200 µL LB medium supplemented with kanamycin and ampicillin in 96-well plates at 37°C with shaking for 5 hours before 3 µL of the cultures were spotted onto LB agar supplemented with 50 μg × mL^−1^ kanamycin, 100 μg × mL^−1^ ampicillin, 0.5 mM IPTG and 40 μg × mL^−1^ X-Gal (5-Bromo-4-chloro-3-indolyl β-D-galactopyranoside). The plates were incubated aerobically at 30 °C overnight. Plates were inspected and photographed after 18 hours and 42 hours of incubation.

### Cell wall isolation and stem peptide analysis

Bacteria from a 1 L culture at OD_550nm_ = 0.4 – 0.5 were harvested at 10 000 *g* for 10 minutes and resuspended in 40 mL of ice-cold 50 mM Tris-HCl, pH 7.0. Cell walls were isolated according to the protocol of Vollmer (2007). Stem peptides were released from the purified cell wall by digestion using LytA as described previously by Straume et al. (2017). Stem peptides from 0.5 mg cell wall were separated by HPLC using a C18 reverse-phase column (Halo 160 Å ES-C18, 2.7 μm, 4.6 x 250 mm from Advanced Material Technology) coupled to a Dionex Ultimate 3000. A 120-minutes linear gradient of acetonitrile from 0% to 15% in 0.05% trifluoroacetic acid with a flow rate of 0.5 mL min^−1^ was used to elute the stem peptides that were detected at 206 nm. The identity of stem peptides was decided by comparison of their retention time with similar chromatograms of mass spectrometry identified peptides published by Gjennestad et al. (2025).

### Statistical analysis

Statistical analyses were conducted in the Rstudio software (Posit, 2026) Version 2026.01.1. Sample results were compared using a one-way ANOVA prior to a Dunnett’s post hoc test to compare each sample to its control or the Wt measurements when appropriate (specified in the figures). Analyses were performed using the functions aov() and glht() from the multcomp package (Hothorn et al., 2008) Version 1.4 - 30.

## Supporting information

Pde1-GFP

CdaA-GFP

## Acknowledgements

The work was funded by the Norwegian University of Life Sciences and a grant from the Research Council of Norway, project nr. 314720. The authors would like to thank Dr. Catherine Sem Wegner and the Advanced Electron Microscopy Core Facility at the Institute for Cancer Research at Oslo University Hospital for help with TEM sample preparation and use of the TEM infrastructure. Thank you to Dr. Vegard Eldholm, Martha Langedok Bjørnstad and Ragnhild Bardal Roness at the Norwegian Institute of Public Health for providing the clinical isolates used in this work.

