## Supplemental figures and tables for "Cyclic di-AMP signalling affects cell division and penicillin susceptibility in *Streptococcus pneumoniae*"

### Supplementary figures

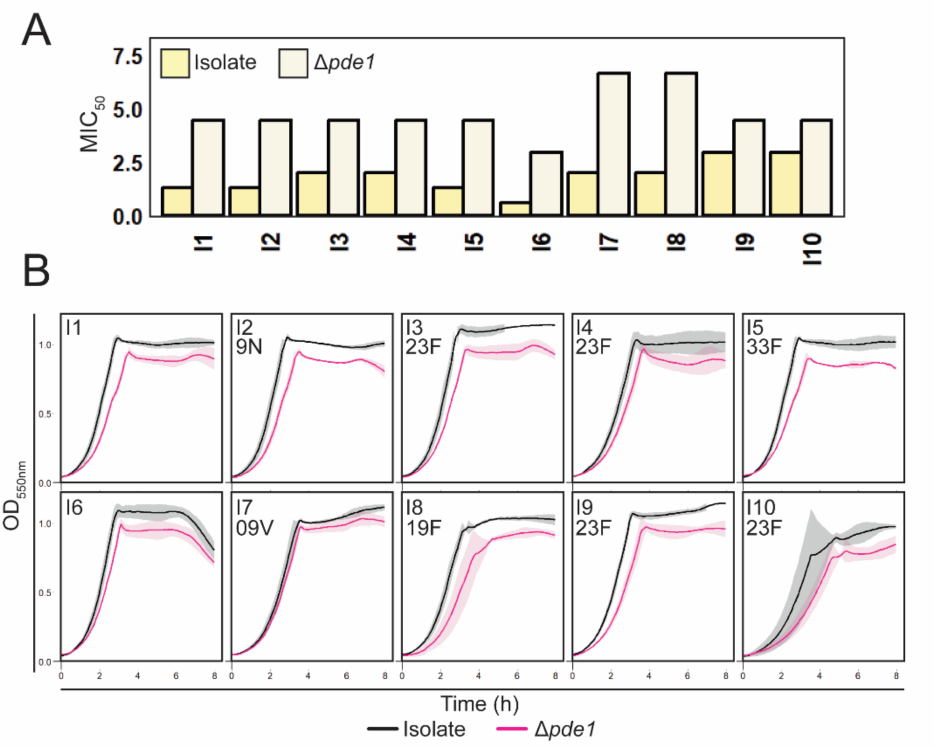


**Figure S1. Penicillin resistance and growth are affected by *pde1* in clinical isolates. (A)** The Penicillin G MIC was measured using broth microdilution for ten clinical isolates (I1-10) and their corresponding Δ*pde1* mutants. The Δ*pde1* mutants showed increased MIC values for all isolates. **(B)** Growth ($OD_{550nm}$) was measured over time and every isolate except I7 showed reduced growth when *pde1* was deleted. The data represent mean and standard deviation of three biological replicates respectively. The serotype of each isolate (when known) is indicated.

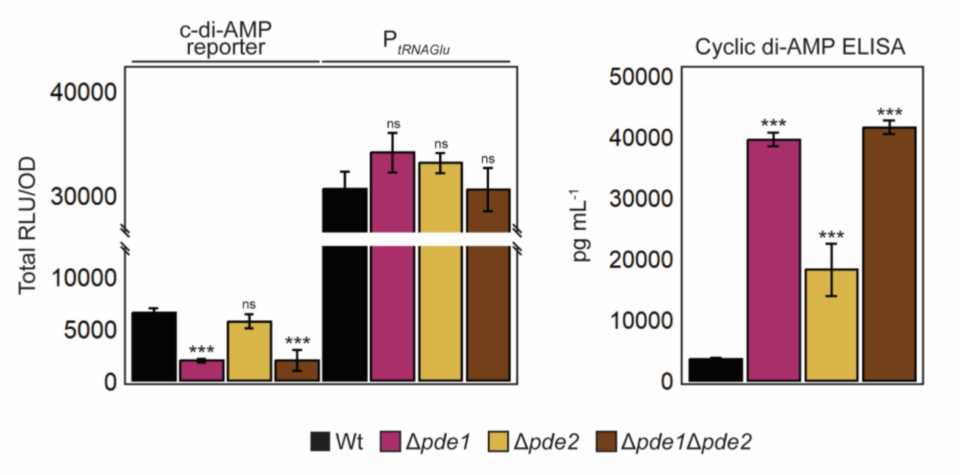


**Figure S2. Comparison of the cyclic di-AMP riboswitch reporter assay and the cyclic di-AMP ELISA.** (left panel) The total luciferase activity (RLU/OD), which is inversely proportional to the cyclic di-AMP levels produced in cells, was measured during 12 hours of growth (c-di-AMP reporter). Luciferase activity in corresponding strains without the riboswitch upstream *luc* was used as control of the $P_{tRNAGlu}$ promoter activity in Wt and mutants. (right panel) Cyclic di-AMP was also measured with a cyclic di-AMP ELISA kit. The results show that both the Δ*pde1* and the double Δ*pde1*Δ*pde2* mutant had significantly higher levels of cyclic di-AMP than the control and that these mutants had approximately the same cyclic di-AMP level. The Δ*pde2* mutant did not show significant difference from the Wt in the *luc* reporter measurement, while the ELISA detected significantly higher cyclic di-AMP levels for this mutant. Each mutant was tested three times and bars, and error bars represent mean and standard deviation respectively. Statistical significance is indicated by *** ($P \leq 0.001$) and ns ($P > 0.1$).

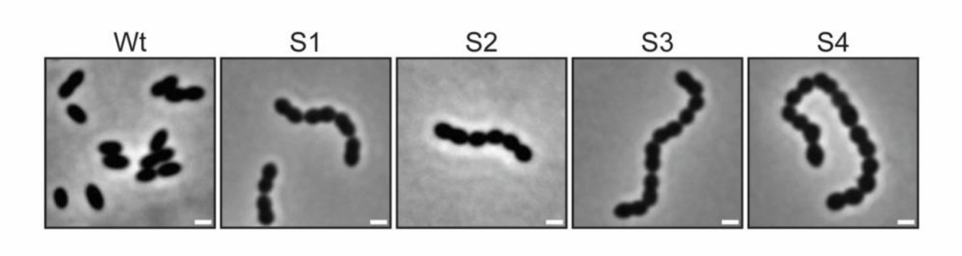


**Figure S3. Morphology of Δ*cdaR* suppressor mutants (S1-4).** The cells appeared smaller and more chained than the Wt, resembling cells with high cyclic di-AMP levels (Δ*pde1* or overexpression of *cdaA*). Whole genome sequencing identified suppressor mutations in *pde1* and *cdaA*.

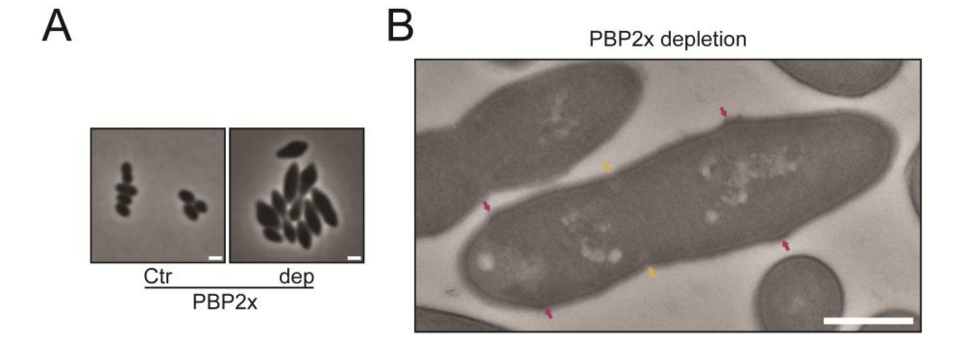


**Figure S4. Morphology of PBP2x depleted *S. pneumoniae*.** (A) Phase contrast microscopy imaging showing the typical lemon-like shape of PBP2x depleted (dep) cells not observed for cells with low cyclic di-AMP levels (*pde1* and *cdaA* overexpression). (B) Transmission electron microscopy imaging showed that PBP2x depleted cells lacked septal cross-walls (yellow arrows) and failed to initiate septal cell wall synthesis at new division sites (likely new division sites are indicated by red arrows). Cells with low cyclic di-AMP levels contained multiple sites of partly produced septal cross walls (Figure 3 in main text). Scale bar is 1 µm and 500 nm for phase contrast and TEM, respectively.

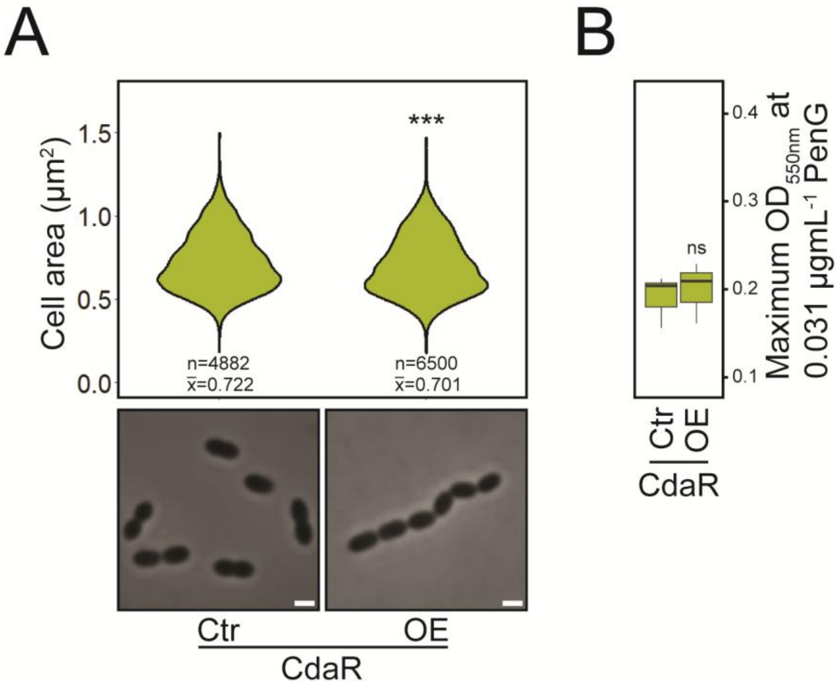


**Figure S5. CdaR overexpression in *S. pneumoniae*.** (A) Violon plots (upper) and phase contrast imaging (lower) showing that overexpression of CdaR marginally reduced (4%) average cell size. (B) Overexpression of CdaR did not influence the susceptibility to Penicillin G (data from three biological replicates). Phase contrast microscopy was performed three times with similar results. Statistical significance: *** (P≤0,001) and ns (P>0.1).

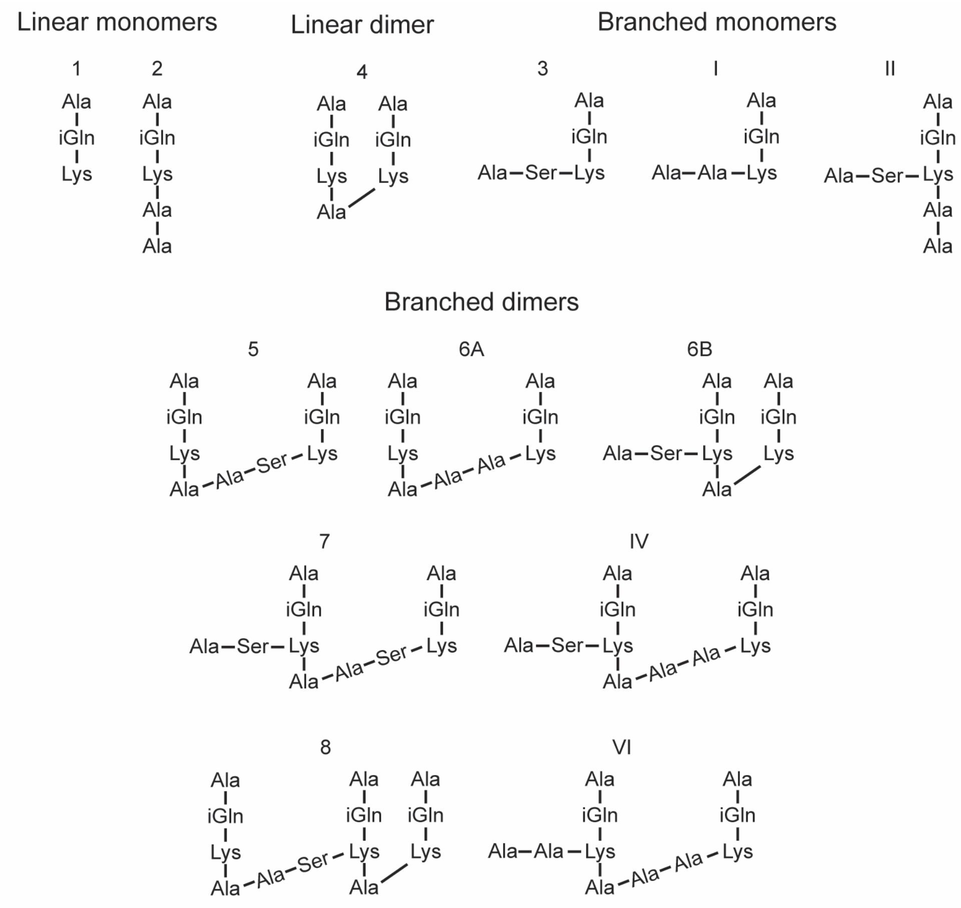


**Figure S6**. Structure of stem peptides.

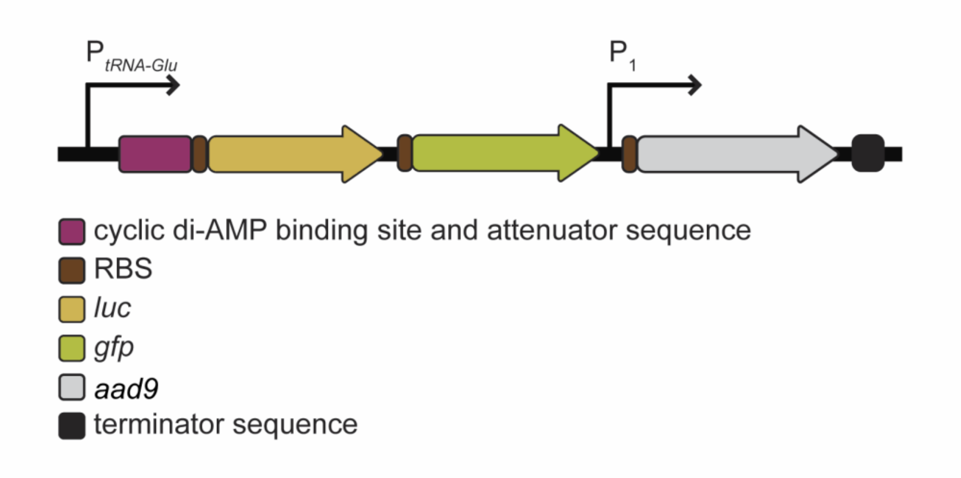


**Figure S7. Organization of the cyclic di-AMP riboswitch reporter.** Transcription of *luc* (luciferase), *gfp* and *aad9* (spectinomycin resistance) was driven by $P_{tRNA\text{-}Glu}$. In addition, *aad9* contained its own constitutive promoter (P1). The riboswitch upstream *kimA* of *Bacillus subtilis* was placed just upstream the ribosomal binding site (RBS) of *luc*. A transcriptional terminator sequence was placed downstream of *aad9*.

### Supplementary Tables

**Table S1: Mutations in suppressor mutants of Δ*cdaR* depleted cells**

| Gene | Locus | DNA difference | Protein difference | Mutant |
|---|---|---|---|---|
| *cdaA* | spr1419 | c.250C-->T | p.84G-->R | S1 |
| *bguD* | spr1834 | c.976G-->T | p326G-->* | S1, S3 |
| *pde1* | spr2010 | c.4delT | p.fsX*6 | S3 |
| | | c.578G-->A | p.193S-->L | S4 |
| | | c.1043A-->T | p.348L-->* | S2 |

* Indicates stop codon
fsX indicates frameshift

**Table S2: Strains and plasmids used in this work**

| Name | Relevant characteristics | Reference |
|---|---|---|
| ***S. pneumoniae* strains:** | | |
| RH425 | R6 derivative, but Δ*comA*::*ermAM*, *rpsL1* $Ery^r$ , $Sm^r$ | (Johnsborg & Håvarstein, 2009) |
| SPH131 | RH425, but Δ*IS1167*::*P1-comR*, Δ*cps*::$P_{comX}$-janus; $Ery^r$, $Kan^r$ | (Berg et al., 2011) |
| SPH155 | Δ*comA* P1::P*comR*::*comR* P*comX*$^c$::*pbp2b* $Sm^r$ | (Berg et al., 2013) |
| SPH162 | Δ*comA* $P_1$::$P_{comR}$::*comR* $P_{comX}$::*pbp2x* $Ery^r$ $Sm^r$ | (Berg et al., 2013) |
| SPH164 | Δ*comA* $P_1$::$P_{comR}$::*comR* $P_{comX}$::*pbp2x* Δ$pbp2x_{Wt}$ $Ery^r$ $Sm^r$ | (Berg et al., 2013) |
| SHA7 | RH425 but Δ*pde1*::janus; $Kan^r$ | This work |
| SHA14 | RH425 but Δ*pde1*::DEL; $Sm^r$ | This work |
| SHA16 | RH425, but Δ*IS1167*::P1-*comR*, Δ*cps*::$P_{comX}$-pde1; $Ery^r$, $Sm^r$ | This work |
| SHA18 | RH425 but Δ*pde2*::janus; $Kan^r$ | This work |
| SHA22 | RH425 but Δ*pde1*::DEL, Δ*pde2*::janus, $Kan^r$ | This work |
| RH425 | R6 derivative, but Δ*comA*::*ermAM*, *rpsL1*; $Ery^r$, $Sm^r$ | (Johnsborg & Håvarstein, 2009) |
| RSG1 | Isolate 1 | This work |
| RSG2 | Isolate 2 | This work |
| RSG3 | Isolate 3 | This work |
| RSG5 | Isolate 4 | This work |

| | | |
|---|---|---|
| RSG7 | Isolate 5 | This work |
| RSG9 | Isolate 6 | This work |
| RSG 48 | Mutant M1 | This work |
| RSG 49 | Mutant M2 | This work |
| RSG 50 | Mutant M3 | This work |
| RSG 51 | Mutant M4 | This work |
| RSG 52 | Mutant M5 | This work |
| RSG210 | Isolate 3 but Δ*pde1*::janus; $Kan^r$ | This work |
| RSG212 | Isolate 6 but Δ*pde1*::janus; $Kan^r$ | This work |
| RSG226 | Isolate 7 | This work |
| RSG228 | Isolate 8 | This work |
| RSG229 | Isolate 9 | This work |
| RSG232 | Isolate 10 | This work |
| RSG245 | Isolate 9 but Δ*pde1*::janus; $Kan^r$ | This work |
| RSG254 | Isolate 1 but Δ*pde1*::janus; $Kan^r$ | This work |
| RSG255 | Isolate 2 but Δ*pde1*::janus; $Kan^r$ | This work |
| RSG256 | Isolate 4 but Δ*pde1*::janus; $Kan^r$ | This work |
| RSG257 | Isolate 5 but Δ*pde1*::janus; $Kan^r$ | This work |
| RSG258 | Isolate 7 but Δ*pde1*::janus; $Kan^r$ | This work |
| RSG259 | Isolate 10 but Δ*pde1*::janus; $Kan^r$ | This work |
| RSG260 | Isolate 8 but Δ*pde1*::janus; $Kan^r$ | This work |
| RSG369 | RH425 but *pde1-sfGFP*; $Sm^r$ | This work |
| RSG439 | RH425, but Δ*IS1167*::*P1-comR*, Δ*cps*::$P_{comX}$-pde2; $Ery^r$, $Sm^r$ | This work |
| RSG440 | RH425, but Δ*IS1167*::*P1-comR*, Δ*cps*::$P_{comX}$-cdaA; $Ery^r$, $Sm^r$ | This work |
| RSG441 | RH425, but Δ*IS1167*::*P1-comR*, Δ*cps*::$P_{comX}$-cdaR; $Ery^r$, $Sm^r$ | This work |
| RSG459 | RH425, but Δ*IS1167*::*P1-comR*, Δ*cps*::$P_{comX}$-*cdaA-linker-sfGFP*; $Ery^r$, $Sm^r$ | This work |
| RSG465 | RH425, but Δ*IS1167*::*P1-comR*, Δ*cps*::$P_{comX}$-*cdaA*, cyclic di-AMP reporter in *zip*-locus; $Ery^r$, $Sm^r$, $Spec^r$ | This work |
| RSG466 | RH425, but Δ*IS1167*::*P1-comR*, Δ*cps*::$P_{comX}^c$-*cdaA*; $Ery^r$, $Sm^r$ | This work |
| RSG469 | RH425, but Δ*IS1167*::*P1-comR*, Δ*cps*::$P_{comX}^c$-*cdaA*, Δ*cdaA*::janus; $Ery^r$, $Kan^r$ | This work |
| RSG474 | RH425, but Δ*IS1167*::*P1-comR*, Δ*cps*::$P_{comX}^c$-*cdaR*; $Ery^r$, $Sm^r$ | This work |
| RSG477 | RH425, but Δ*IS1167*::*P1-comR*, Δ*cps*::$P_{comX}$-*GFP-linker-cdaR*; $Ery^r$, $Sm^r$ | This work |

| | | |
|---|---|---|
| RSG481 | RH425, but Δ*IS1167*::*P1-comR*, Δ*cps*::$P_{comX}$-*pde1*, cyclic di-AMP reporter in *zip*-locus; Ery$^r$, Sm$^r$, Spec$^r$ | This work |
| RSG482 | RH425 but cyclic di-AMP reporter in *zip*-locus; Sm$^r$, Spec$^r$ | This work |
| RSG484 | RH425, but Δ*IS1167*::*P1-comR*, Δ*cps*::$P_{comX}^{c}$-*cdaR*, Δ*cdaR*::janus; Ery$^r$, Kan$^r$ | This work |
| RSG486 | RH425 but Δ*pde1*::DEL, cyclic di-AMP reporter in *zip*-locus; Sm$^r$, Spec$^r$ | This work |
| RSG490 | RH425 but $tRNA_{Glu}$ reporter in *zip*-locus; Sm$^r$, Spec$^r$ | This work |
| RSG494 | RH425 but Δ*pde1*::DEL, $tRNA_{Glu}$ reporter in *zip*-locus; Sm$^r$, Spec$^r$ | This work |
| RSG495 | RH425 but Δ*pde1*::DEL, Δ*pde2*::janus, cyclic di-AMP reporter in *zip*-locus; Sm$^r$, Spec$^r$ | This work |
| RSG496 | RH425 but Δ*pde2*::janus, cyclic di-AMP reporter in *zip*-locus; Kan$^r$, Spec$^r$ | This work |
| RSG497 | RH425 but Δ*pde2*::janus, $tRNA_{Glu}$ reporter in *zip*-locus; Kan$^r$, Spec$^r$ | |
| RSG498 | Suppressor mutant S1 | This work |
| RSG499 | Suppressor mutant S2 | This work |
| RSG500 | Suppressor mutant S3 | This work |
| RSG501 | Suppressor mutant S4 | This work |
| RSG514 | RH425, but Δ*IS1167*::*P1-comR*, Δ*cps*::$P_{comX}$-pde2, $tRNA_{Glu}$ reporter in *zip*-locus; Ery$^r$, Sm$^r$ | This work |
| RSG519 | RH425 but Δ*pde1*::DEL, Δ*pde2*::janus, $tRNA_{Glu}$ reporter in *zip*-locus; Sm$^r$, Spec$^r$ | This work |
| RSG521 | RH425, but Δ*IS1167*::*P1-comR*, Δ*cps*::$P_{comX}$-*pde1*, $tRNA_{Glu}$ reporter in *zip*-locus; Ery$^r$, Sm$^r$, Spec$^r$ | This work |
| RSG526 | RH425, but Δ*IS1167*::*P1-comR*, Δ*cps*::$P_{comX}$-*cdaA*, $tRNA_{Glu}$ reporter in *zip*-locus; Ery$^r$, Sm$^r$, Spec$^r$ | This work |
| RSG533 | RH425 but *pde2-sfGFP*; Sm$^r$ | This work |
| ***Streptococcus oralis* strains:** | | |
| Uo5 | A high level β-lactam resistant isolate of *S. oralis* isolated from a Hungarian nasal swab | (Reichmann et al., 1997) |
| ***Escherichia coli* strains:** | | |
| XL-1 Blue | Host strain | Agilent Technologies |
| BTH101 | BACTH expression strain, *cya-* | Euromedex |
| **Plasmids:** | | |

| | | |
|---|---|---|
| pUT18 | Plasmid used in BACTH analysis | Euromedex |
| pUT18C | Plasmid used in BACTH analysis | Euromedex |
| pKT25 | Plasmid used in BACTH analysis | Euromedex |
| pKNT25 | Plasmid used in BACTH analysis | Euromedex |
| pKT25-zip | T25 fused to a leucine zipper domain | Euromedex |
| pUT18C-zip | T18 fused to a leucine zipper domain | Euromedex |
| pUT18-pde1 | T18 domain fused to the C-terminus of Pde1 | This work |
| pKNT25-pde1 | T25 domain fused to the C-terminus of Pde1 | This work |
| pKNT25-pde2 | T25 domain fused to the C-terminus of Pde2 | This work |
| pUT18-pde2 | T18 domain fused to the C-terminus of Pde2 | This work |
| pKNT25-cdaA | T25 domain fused to the C-terminus of CdaA | This work |
| pUT18-cdaA | T18 domain fused to the C-terminus of CdaA | This work |
| pUT18C-cdaR | T18 domain fused to the N-terminus of CdaR | This work |
| pKT25-cdaR | T25 domain fused to the N-terminus of CdaR | This work |

**Table S3: Primers used in this work**

| Primer | Description | Sequence 5'-->3' | Reference |
|---|---|---|---|
| **Primers used to amplify the Janus cassette** | | | |
| Janus F | Start Janus cassette | GTTTGATTTTTAATGGATAATGTG | (Johnsborg et al., 2008) |
| Janus R | End Janus cassette | CTTTCCTTATGCTTTTGGAC | (Johnsborg et al., 2008) |
| **Primers to create the Δ*pde1*::janus amplicon and sequence the *pde1* gene** | | | |
| KHB94 | Janus R, **overlap revers compliment P1 promoter** | **AAGTATTTTCTAGTATTATAGCACATTTAA**CTTTCCTTATGCTTTTGGAC | This work |
| SHA8 | Last 78bp in *pde1* **overlap P1 promoter** | **GTGCTATAATACTAGAAAATACTT**GTAACCTTGTCAGAAGCAGG | This work |
| SHA5 | Start Janus cassette, **overlap just up *pde1*** | CACATTATCCATTAAAAATCAAAC**TCAAAACCTCTTGGCACCCA** | This work |
| SHA4 | ~1000bp upstream *pde1* | GAGAAAAGAGAAGAGAGAGCC | This work |
| SHA7 | ~1000bp downstream *pde1* | GTTGAGATTCAACACATCTCG | This work |
| SHA1 | ~150bp downstream *pde1* | CAGCGTGAGCTTTTTCTTCC | This work |
| SHA2 | ~650bp in *pde1* | CATGATGTTTTCTCGTCGGG | This work |
| SHA3 | ~150bp upstream *pde1* | GCGAAATTGGTTTGTTAGAGG | This work |
| **Primers to create the Δ*pde1*::DEL amplicon** | | | |
| SHA15 | P1 promoter (from Δpde1::janus amplicon) **overlap revers just up *pde1*** | **TGGGTGCCAAGAGGTTTTGA**TTAAATGTGCTATAATACTAGAAAA | This work |
| SHA4 | ~1000bp upstream *pde1* | GAGAAAAGAGAAGAGAGAGCC | This work |
| SHA7 | ~1000bp downstream *pde1* | GTTGAGATTCAACACATCTCG | This work |
| **Primers to create the Δ*pde2*::janus amplicon and sequence the *pde2* gene** | | | |
| SHA11 | ~1000bp upstream *pde2* | CAGCTAGAAAAATTCCTAGCG | This work |
| SHA12 | Start of Janus cassette, **overlap just up *pde2*** | CACATTATCCATTAAAAATCAAAC**AAATAACCTCAATTCTTTCTCATTT** | This work |

| | | | |
|---|---|---|---|
| SHA13 | End of Janus cassette, **overlap just down *pde2*** | GTCCAAAAGCATAAGGAAAG**TAAAATACTTGCCAAACTTTTCAG** | This work |
| SHA14 | ~1000bp downstream *pde2* | TGGTCAGAAATTATTGGAAGGC | This work |
| RSG133 | ~150bp upstream *pde2* | ATTACCGGTCATACTGGC | This work |
| RSG134 | ~150bp downstream *pde2* | CCATGAAGACAACTGGGCG | This work |
| RSG135 | ~800bp in *pde2* | GAGCATGATGGTGGAGGCC | This work |
| **Primers to amplify the $P_{comX}$::janus amplicon and create and replacements** | | | |
| KHB31 | ~800bp upstream $P_{comX}$ | ATAACAAATCCAGTAGCTTTGG | (Berg et al., 2011) |
| KHB34 | ~800bp downstream $P_{comX}$::janus | CATCGGAACCTATACTCTTTTAG | (Berg et al., 2011) |
| KHB33 | just down $P_{comX}$::janus | TTTCTAATATGTAACTCTTCCCAAT | (Berg et al., 2011) |
| KHB36 | end of $P_{comX}$ | TGAACCTCCAATAATAAATATAAAT | (Berg et al., 2011) |
| | Template strain: SPH131 (RH425, but Δ*is1167*::*p1-comR*, Δ*cps*::$P_{comX}$::janus) | | (Berg et al., 2011) |
| **Primers to amplify the $P_{comX}^{c}$::janus amplicon** | | | |
| KHB31 | ~800bp upstream $P_{comX}$ | ATAACAAATCCAGTAGCTTTGG | (Berg et al., 2011) |
| KHB34 | ~800bp downstream $P_{comX}$::janus | CATCGGAACCTATACTCTTTTAG | (Berg et al., 2011) |
| KHB33 | just down $P_{comX}$::janus | TTTCTAATATGTAACTCTTCCCAAT | (Berg et al., 2011) |
| KHB147 | end of $P_{comX}$ with **SNP** | TGAACCTC**G**AATAATAAATATAAATTCTGTAATTAG | (Berg et al., 2013) |
| | Template strain: SPH155 (RH425, but Δ*comA* p1::$P_{comR}$::*comR* $P_{comX}^{c}$::*pbp2b* $sm^r$) | | (Berg et al., 2013) |

| | | | |
|---|---|---|---|
| **Primers to create the $P_{comX}$::*pde1* amplicon** | | | |
| KHB31 | ~800bp upstream $P_{comX}$ | ATAACAAATCCAGTAGCTTTGG | (Berg et al., 2011) |
| KHB34 | ~800bp downstream $P_{comX}$::janus | CATCGGAACCTATACTCTTTTAG | (Berg et al., 2011) |
| SHA9 | End of $P_{comX}$, **overlap start *pde1*** | ATTTATATTTATTATTGGAGGTTCA**ATGAAAAAATTTTATGTAAGTCCAA** | This work |
| SHA10 | Just down $P_{comX}$::janus, **overlap end *pde1*** | ATTGGGAAGAGTTACATATTAGAAA**TCATTCTTCTTTCTCCTTTTCC** | This work |
| **Primers to create the $P_{comX}$::*cdaA* amplicon** | | | |
| KHB31 | ~800bp upstream $P_{comX}$ | ATAACAAATCCAGTAGCTTTGG | (Berg et al., 2011) |
| KHB34 | ~800bp downstream $P_{comX}$::janus | CATCGGAACCTATACTCTTTTAG | (Berg et al., 2011) |
| RSG220 | End of $P_{comX}$, **overlap start *cdaA*** | ATTTATATTTATTATTGGAGGTTCA**ATGAATTTTCAACAATTATCCAATC** | This work |
| RSG221 | Just down $P_{comX}$::janus, **overlap end *cdaA*** | ATTGGGAAGAGTTACATATTAGAAA**CTATTTTTTTCATGTTTCCATCC** | This work |
| **Primers to create the $P_{comX^c}$::*cdaA* amplicon** | | | |
| KHB31 | ~800bp upstream $P_{comX}$ | ATAACAAATCCAGTAGCTTTGG | (Berg et al., 2011) |
| KHB34 | ~800bp downstream $P_{comX}$::janus | CATCGGAACCTATACTCTTTTAG | (Berg et al., 2011) |
| RSG270 | End of $P_{comX^c}$, **overlap start *cdaA*** | ATTTATATTTATTATTCGAGGTTCA**ATGAATTTTCAACAATTATCCAATC** | This work |
| RSG221 | Just down $P_{comX}$::janus, **overlap end *cdaA*** | ATTGGGAAGAGTTACATATTAGAAA**CTATTTTTTTCATGTTTCCATCC** | This work |
| **Primers to create the Δ*cdaA*::janus amplicon** | | | |

| | | | |
|---|---|---|---|
| RSG222 | ~1000bp upstream *cdaA*, ~2000bp upstream *cdaR* | AGCGTTGTTGACAGCAGTTG | This work |
| RSG223 | ~1000bp downstream *cdaA*, ~200bp downstream *cdaR* | GACCTGCCACCAAGGCCG | This work |
| RSG224 | Start of Janus cassette, **overlap 72bp in *cdaA*** | CACATTATCCATTAAAAATCAAAC**GATAGCTATCGTCCATGGAC** | This work |
| RSG225 | End of Janus cassette, **overlap 732bp in (keep the last 126bp) *cdaA*** | GTCCAAAAGCATAAGGAAAG**TTTAAGCACAACCTAACACTTG** | This work |
| **Primers to create the $P_{comX}$::*cdaR* amplicon** | | | |
| KHB31 | ~800bp upstream $P_{comX}$ | ATAACAAATCCAGTAGCTTTGG | (Berg et al., 2011) |
| KHB34 | ~800bp downstream $P_{comX}$::janus | CATCGGAACCTATACTCTTTTAG | (Berg et al., 2011) |
| RSG228 | End of $P_{comX}$, **overlap start *cdaR*** | ATTTATATTTATTATTGGAGGTTCA**ATGAAAAAAATAGTTTATATATCATA** | This work |
| RSG229 | Just down $P_{comX}$::janus, **overlap end *cdaR*** | ATTGGGAAGAGTTACATATTAGAAA**TTAATTCGTTTTTGAACTAGTTGC** | This work |
| **Primers to create the $P_{comX}^{c}$::*cdaR* amplicon** | | | |
| KHB31 | ~800bp upstream $P_{comX}$ | ATAACAAATCCAGTAGCTTTGG | (Berg et al., 2011) |
| KHB34 | ~800bp downstream $P_{comX}$::janus | CATCGGAACCTATACTCTTTTAG | (Berg et al., 2011) |
| RSG269 | End of $P_{comX}^{c}$, **overlap start *cdaR*** | ATTTATATTTATTATTCGAGGTTCA**ATGAAAAAAATAGTTTATATATCATA** | This work |
| RSG229 | Just down $P_{comX}$::janus, **overlap end *cdaR*** | ATTGGGAAGAGTTACATATTAGAAA**TTAATTCGTTTTTGAACTAGTTGC** | This work |
| **Primers to create the Δ*cdaR*::janus amplicon** | | | |
| RSG222 | ~1000bp upstream *cdaA*, ~2000bp upstream *cdaR* | AGCGTTGTTGACAGCAGTTG | This work |
| RSG230 | ~1000bp downstream *cdaR* | GACCAGACTGTTCACCACC | This work |

| | | | |
|---|---|---|---|
| RSG231 | Start of Janus cassette, **overlap 72bp in *cdaR*** | CACATTATCCATTAAAAATCAAAC**CGCCGTAGCATAGACAAATAA** | This work |
| RSG232 | End of Janus cassette, **overlap 732bp in (keep the last 48bp) *cdaR*** | GTCCAAAAGCATAAGGAAAG**TCGGAGACATCTTCGTCAAC** | This work |
| **Primers to create the $P_{comX}$::*pde2* amplicon** | | | |
| KHB31 | ~800bp upstream $P_{comX}$ | ATAACAAATCCAGTAGCTTTGG | (Berg et al., 2011) |
| KHB34 | ~800bp downstream $P_{comX}$::janus | CATCGGAACCTATACTCTTTTAG | (Berg et al., 2011) |
| RSG196 | End of $P_{comX}$, **overlap start *pde2*** | ATTTATATTTATTATTGGAGGTTCA**ATGGAGATTTGCCAACAAATTTT** | This work |
| RSG197 | Just down $P_{comX}$::janus, **overlap end *pde2*** | ATTGGGAAGAGTTACATATTAGAAA**TCAGTTTTTAAGCAAGTTTTTAAC** | This work |
| **Primers to amplify *sfGFP*** | | | |
| DS323 | Forward *sfGFP* (without start-codon) | AAACATCTTACCGGTTCTAAAG | This work |
| DS233 | Revers *sfGFP* | TTATGCGGCCGCTCCACTAG | This work |
| ATS4 | Revers *sfGFP* (without stop-codon) | TGCGGCCGCTCCACTAGTTTTG | (Mårli et al., 2025) |
| | Template strain: SPH470 (Δ*comA, m(sf)GFP-mltG*; $Ery^R$, $Sm^R$) | | (Stamsås et al., 2017) |
| **Primers to create the *pde1-sfGFP* amplicon** | | | |
| SHA4 | ~1000bp upstream *pde1* | GAGAAAAGAGAAGAGAGAGCC | This work |
| SHA7 | ~1000bp downstream *pde1* | GTTGAGATTCAACACATCTCG | This work |
| RSG126 | End of *sfGFP*, **overlap 2968bp in (keep the last 17bp) of *pde1*** | CTAGTGGAGCGGCCGCATAA**AGGAGAAAGAAGAATGAAAGT** | This work |
| RSG116 | Start of *sfGFP*, **overlap end of *pde1*** | CTTTAGAACCGGTAAGATGTTT**TTCTTTCTCCTTTTCCTTTATTTC** | This work |
| **Primers to create the $P_{comX}$::*cdaA-linker-sfGFP* amplicon** | | | |

| | | | |
|---|---|---|---|
| KHB31 | ~800bp upstream $P_{comX}$ | ATAACAAATCCAGTAGCTTTGG | (Berg et al., 2011) |
| KHB34 | ~800bp downstream $P_{comX}$::janus | CATCGGAACCTATACTCTTTTAG | (Berg et al., 2011) |
| RSG227 | Just down $P_{comX}$::janus, **overlap end of *sfGFP*** | ATTGGGAAGAGTTACATATTAGAAA**TTATGCGGCCGCTCCACTAG** | This work |
| RSG234 | Start of *sfGFP*, <u>linker</u> **overlap end of *cdaA*** | CTTTAGAACCGGTAAGATGTTT<u>TCCAGCTTTAGCTGCAGCTTCTCCACCAGATCC</u>**TTTTTTTTCATGTTTCCATCCTC** | This work |
| RSG220 | End of $P_{comX}$, **overlap start *cdaA*** | ATTTATATTTATTATTGGAGGTTCA**ATGAATTTTCAACAATTATCCAATC** | This work |
| **Primers to create the *pde2-linker-sfGFP* amplicon** | | | |
| SHA11 | ~1000bp upstream *pde2* | CAGCTAGAAAAATTCCTAGCG | This work |
| SHA14 | ~1000bp downstream *pde2* | TGGTCAGAAATTATTGGAAGGC | This work |
| RSG297 | Start of *sfGFP*, <u>linker</u> **overlap end of *pde2*** | CTTTAGAACCGGTAAGATGTTT<u>TCCAGCTTTAGCTGCAGCTTCTCCACCAGATCC</u>**GTTTTTAAGCAAGTTTTTTAACTTT** | This work |
| RSG298 | End of *sfGFP*, **overlap just down *pde2*** | CTAGTGGAGCGGCCGCATAA**TAAAATACTTGCCAAACTTTTCAG** | This work |
| **Primers to create the $P_{comX}$::*cdaR-linker-sfGFP* amplicon** | | | |
| KHB31 | ~800bp upstream $P_{comX}$ | ATAACAAATCCAGTAGCTTTGG | (Berg et al., 2011) |
| KHB34 | ~800bp downstream $P_{comX}$::janus | CATCGGAACCTATACTCTTTTAG | (Berg et al., 2011) |
| RSG229 | Just down $P_{comX}$::janus, **overlap end *cdaR*** | ATTGGGAAGAGTTACATATTAGAAA**TTAATTCGTTTTTGAACTAGTTGC** | This work |

| | | | |
|---|---|---|---|
| RSG274 | End of *sfGFP*, linker **overlap start of *cdaR*** | CAAAACTAGTGGAGCGGCCGCAGGATCTGGTGGAGAAGCTGCAGCTAAAGCTGG**AAAAAAAAATAGTTTATATATCATATCC** | This work |
| RSG275 | End of P$_{comX}$, **overlap start of *sfGFP*** | ATTTATATTTATTATTGGAGGTTCA**ATGAAACATCTTACCGGTTCTAAAG** | This work |
| **Primers to amplify *luc-sfGFP*** | | | |
| RSG248 | Forward *luc* with **RBS** upstream | **AGGAGG**AATAATGAGATCCGC | This work |
| RM218 | Revers *sfGFP* | TTACTTATAAAGCTCATCCATGCC | This work |
| | Template strain: DSM2 (Δ*bgaA*::P$_{ssbB}$-*luc*-*gfp*) | | (Moreno-Gamez et al., 2017) |
| **Primers to create the c-di-AMP reporter amplicon** | | | |
| RSG258 | ~1000bp downstream *zip*-locus (spr1750) | CTTACTCAAGATAAGATTGCTG | This work |
| RSG259 | ~1000bp upstream *zip*-locus (spr1750) | GGAGGGAAGAAGAGGTTTCC | This work |
| RSG260 | Start of *tRNA$_{Glu}$* promoter, **overlap spr1750** | ACTCTTACTATTATACAGTCTTTTCAAACTTTGTCAACTACTTTT**CATAGACATTGGAACTGAAGAA** | This work |
| RSG261 | End of *tRNA$_{Glu}$* promoter, **overlap start of riboswitch sequence** | AAGACTGTATAATAGTAAGAGTTGAAAATAACAACTCA**GAAAACAAATCGCTTAATCTGAAA** | This work |
| RSG262 | Start of *luc*, **overlap end of riboswitch sequence** | GCGGATCTCATTATTCCTCCT**CGATGTCTTCCCCTTTTAATTT** | This work |
| RSG263 | End of *GFP*, **overlap start of *aad9* sequence** | GGCATGGATGAGCTTTATAAGTAA**TTAAATGTGCTATAATACTAGAAAAT** | This work |
| RSG264 | End of *aad9*, **overlap terminator sequence** | CTATTTAAATAACAGATTAAAAAAATTATAA**CCAGGCATCAAATAAAACGAAAG** | This work |
| RSG265 | In spr1750 | CTCACAATCTCAGCTGAACC | This work |
| RSG266 | In spr1750, **overlap end of terminator sequence** | GGTTCAGCTGAGATTGTGAG**CGCAGAAAGGCCCACCCG** | This work |

| | | | |
|---|---|---|---|
| | Template strains: the riboswitch sequence was amplified from *Bacillus subtilis* ATCC6051[1], the spectinomycin resistance marker (*aad9*) was amplified from DS789 (Straume et al., 2020), and the terminator sequence was amplified from pPEPY (Keller et al., 2019) | | |
| **Primers to create the P$_{tRNAGlu}$ reporter** | | | |
| RSG258 | ~1000bp downstream *zip*-locus (spr1750) | CTTACTCAAGATAAGATTGCTG | This work |
| RSG259 | ~1000bp upstream *zip*-locus (spr1750) | GGAGGGAAGAAGAGGTTTCC | This work |
| RSG260 | Start of *tRNA$_{Glu}$* promoter, **overlap spr1750** | ACTCTTACTATTATACAGTCTTTTCAAACTTTGTCAACTACTTTT**CATAGACATTGGAACTGAAGAA** | This work |
| RSG294 | End of *tRNA$_{Glu}$* promoter, **overlap start of *luc* sequence** | AAGACTGTATAATAGTAAGAGT**AGGAGGAATAATGAGATCCGC** | This work |
| RSG263 | End of *GFP*, **overlap start of *aad9* sequence** | GGCATGGATGAGCTTTATAAGTAA**TTAAATGTGCTATAATACTAGAAAAT** | This work |
| RSG264 | End of *aad9*, **overlap terminator sequence** | CTATTTAAATAACAGATTAAAAAAATTATAA**CCAGGCATCAAATAAAACGAAAG** | This work |
| RSG265 | In spr1750 | CTCACAATCTCAGCTGAACC | This work |
| RSG266 | In spr1750, **overlap end of terminator sequence** | GGTTCAGCTGAGATTGTGAG**CGCAGAAAGGCCCACCCG** | This work |

[1]The ATCC6051 was a gift from Dr. Annette Fagerlund.

### Supplementary text

**Text S1: Cyclic di-AMP reporter construct**

Colours are as follows:

**$P_{tRNAGlu}$ promoter**

**Cyclic di-AMP binding site and attenuator sequence**

**Ribosomal binding site (RBS)**

***Luciferase* encoding gene**

***gfp* encoding gene**

**aad9 spectinomycin resistance gene**

**Terminator sequence**

>cyclic_di_AMP_reporter_construct

**AAAAGTAGTTGACAAAGTTTGAAAAGACTGTATAATAGTAAGA**GT
TGAAAATAACAACTCA**GAAAACAAATCGCTTAATCTGAAATCAGAG**
**CGGGGGACCCAATAGAACGGCTTTTTGCCGTTGGGGTGAATCCTT**
**TTTAGGTAGGGCTAACTCTCATATGCCCGAATCCGTCAGCTAACC**
**TCGTAAGCGTTCGTGAGAGGAGATGAATGAAACCTGTGTTCGATG**
**TTATGGCACAGGGGCATCCGTTTGCCTCTGTGTTTTTTGTTGTTC**
**ATTTTTGAATGCAAATCGCATACAGAGACATCATAGCAGAAAGGA**
**CTATCATCAACATGTGAGAGCAAGACACTGATGGATGTTTTATTG**
**TTTTAATTGAAAAATGAGAAATTAAAAGGGGAAGACATCGAGGAG**
**G**AATA**ATGAGATCCGCCAAAAACATAAAGAAAGGCCCGGCGCCAT**
**TCTATCCTCTAGAGGATGGAACCGCTGGAGAGCAACTGCATAAGG**
**CTATGAAGAGATACGCCCTGGTTCCTGGAACAATTGCTTTTACAG**
**ATGCACATATCGAGGTGAACATCACGTACGCGGAATACTTCGAAA**
**TGTCCGTTCGGTTGGCAGAAGCTATGAAACGATATGGGCTGAATA**
**CAAATCACAGAATCGTCGTATGCAGTGAAAACTCTCTTCAATTCT**
**TTATGCCGGTGTTGGGCGCGTTATTTATCGGAGTTGCAGTTGCGC**
**CCGCGAACGACATTTATAATGAACGTGAATTGCTCAACAGTATGA**
**ACATTTCGCAGCCTACCGTAGTGTTTGTTTCCAAAAAGGGGTTGC**
**AAAAAATTTTGAACGTGCAAAAAAAATTACCAATAATCCAGAAAA**
**TTATTATCATGGATTCTAAAACGGATTACCAGGGATTTCAGTCGA**
**TGTACACGTTCGTCACATCTCATCTACCTCCCGGTTTTAATGAATA**
**CGATTTTGTACCAGAGTCCTTTGATCGTGACAAAACAATTGCACT**

**GATAATGAATTCCTCTGGATCTACTGGGTTACCTAAGGGTGTGGC
CCTTCCGCATAGAACTGCCTGCGTCAGATTCTCGCATGCCAGAGA
TCCTATTTTTGGCAATCAAATCATTCCGGATACTGCGATTTTAAGT
GTTGTTCCATTCCATCACGGTTTTGGAATGTTTACTACACTCGGAT
ATTTGATATGTGGATTTCGAGTCGTCTTAATGTATAGATTTGAAG
AAGAGCTGTTTTTACGATCCCTTCAGGATTACAAAATTCAAAGTG
CGTTGCTAGTACCAACCCTATTTTCATTCTTCGCCAAAAGCACTCT
GATTGACAAATACGATTTATCTAATTTACACGAAATTGCTTCTGG
GGGCGCACCTCTTTCGAAAGAAGTCGGGGAAGCGGTTGCAAAAC
GCTTCCATCTTCCAGGGATACGACAAGGATATGGGCTCACTGAGA
CTACATCAGCTATTCTGATTACACCCGAGGGGGATGATAAACCGG
GCGCGGTCGGTAAAGTTGTTCCATTTTTTGAAGCGAAGGTTGTGG
ATCTGGATACCGGGAAAACGCTGGGCGTTAATCAGAGAGGCGAA
TTATGTGTCAGAGGACCTATGATTATGTCCGGTTATGTAAACAAT
CCGGAAGCGACCAACGCCTTGATTGACAAGGATGGATGGCTACA
TTCTGGAGACATAGCTTACTGGGACGAAGACGAACACTTCTTCAT
AGTTGACCGCTTGAAGTCTTTAATTAAATACAAAGGATATCAGGT
GGCCCCCGCTGAATTGGAATCGATATTGTTACAACACCCCAACAT
CTTCGACGCGGGCGTGGCAGGTCTTCCCGACGATGACGCCGGTG
AACTTCCCGCCGCCGTTGTTGTTTTGGAGCACGGAAAGACGATGA
CGGAAAAAGAGATCGTGGATTACGTCGCCAGTCAAGTAACAACC
GCGAAAAAGTTGCGCGGAGGAGTTGTGTTTGTGGACGAAGTACC
GAAAGGTCTTACCGGAAAACTCGACGCAAGAAAAATCAGAGAGA
TCCTCATAAAGGCCAAGAAGGGCGGAAAGCCCAAATTGTAA**GATC
CACTAGATACTGATTAACTAATA**AGGAGG**ACAAACATGTCAAAAGG
AGAAGAGCTGTTCACAGGTGTTGTGCCGATTCTCGTTGAGCTTGACGG
AGATGTAAACGGACACAAATTCTCTGTTCGCGGTGAAGGTGAAGGAG
ATGCAACAAACGGCAAGCTGACATTGAAGTTTATTTGCACAACTGGA
AAGCTGCCGGTTCCTTGGCCGACACTTGTAACGACGCTGACTTACGGC
GTTCAATGCTTCTCTCGTTATCCAGACCACATGAAACGCCATGATTTC
TTCAAATCTGCAATGCCTGAAGGCTACGTTCAAGAGCGTACGATCAG
CTTCAAAGATGACGGAACGTACAAAACAAGAGCAGAAGTGAAGTTTG
AAGGTGACACACTTGTGAACCGCATTGAATTGAAAGGCATTGATTTC
AAAGAAGATGGAAACATCCTTGGACACAAACTTGAATACAACTTCAA

CAGCCACAACGTATACATCACTGCTGACAAACAAAAAAACGGCATCA
AAGCAAACTTCAAAATCCGTCATAACGTAGAGGACGGTTCTGTTCAG
CTTGCTGATCATTATCAGCAAAATACACCGATCGGTGACGGCCCGGTT
CTTCTTCCTGATAACCATTATTTATCAACTCAAAGCGTATTATCAAAA
GACCCAAATGAAAAGCGTGACCACATGGTGCTGCTTGAATTTGTGAC
AGCTGCTGGTATCACTCACGGCATGGATGAGCTTTATAAGTAATTAAA
TGTGCTATAATACTAGAAAATACTTGT**GGAGG**TTCCATT**GTGAGGAG**
**GATATATTTGAATACATACGAACAAATTAATAAAGTGAAAAAAAT**
**ACTTCGGAAACATTTAAAAAATAACCTTATTGGTACTTACATGTTT**
**GGATCAGGAGTTGAGAGTGGTCTAAAACCAAATAGTGATCTTGAC**
**TTTTTAGTCGTCGTATCTGAACCATTGACAGATCAAAGTAAAGAA**
**ATACTTATACAAAAAATTAGACCTATTTCAAAAAAAAATAGGAGAT**
**AAAAGCAACTTACGATATATTGAATTAACAATTATTATTCAGCAAG**
**AAATGGTACCGTGGAATCATCCTCCCAAACAAGAATTTATTTATG**
**GAGAATGGTTACAAGAGCTTTATGAACAAGGATACATTCCTCAGA**
**AGGAATTAAATTCAGATTTAACCATAATGCTTTACCAAGCAAAAC**
**GAAAAAATAAAAGAATATACGGAAATTATGACTTAGAGGAATTAC**
**TACCTGATATTCCATTTTCTGATGTGAGAAGAGCCATTATGGATT**
**CGTCAGAGGAATTAATAGATAATTATCAGGATGATGAAACCAACT**
**CTATATTAACTTTATGCCGTATGATTTTAACTATGGACACGGGTAA**
**AATCATACCAAAAGATATTGCGGGAAATGCAGTGGCTGAATCTTC**
**TCCATTAGAACATAGGGAGAGAATTTTGTTAGCAGTTCGTAGTTA**
**TCTTGGAGAGAATATTGAATGGACTAATGAAAATGTAAATTTAAC**
TATAAACTATTTAAATAACAGATTAAAAAAATTATAA**CCAGGCATC**
**AAATAAAACGAAAGGCTCAGTCGAAAGACTGGGCCTTTCGTTTTA**
**TCTGTTGTTTGTCGGTGAATGCTCTCTACTAGAGTCACACTGGCT**
**CACCTTCGGGTGGGCCTTTCTGCG**
